# Daily glycome and transcriptome profiling reveals polysaccharide structures and glycosyltransferases critical for cotton fiber growth

**DOI:** 10.1101/2024.04.23.589927

**Authors:** Sivakumar Swaminathan, Corrinne E. Grover, Alither S. Mugisha, Lauren E. Sichterman, Youngwoo Lee, Pengcheng Yang, Eileen L. Mallery, Josef J Jareczek, Alexis G Leach, Jun Xie, Jonathan F. Wendel, Daniel B. Szymanski, Olga A. Zabotina

**Affiliations:** Roy J Carver Department of Biochemistry, Biophysics and Molecular Biology, Iowa State University, Ames, IA 50011, USA; Department of Ecology, Evolution and Organismal Biology, Iowa State University, Ames, IA 50011, USA; Department of Biological Sciences, Purdue University, West Lafayette, IN 47907, USA; Department of Statistics, Purdue University, West Lafayette, IN 47907, USA

## Abstract

Cotton fiber length and strength are key determinants of its quality. Dynamic changes in the pectin, xyloglucan, xylan, and cellulose polysaccharide epitopes content during fiber growth contribute to complex remodeling of fiber cell wall (CW) and quality. Detailed knowledge about polysaccharide compositional and structural alteration in the fiber during fiber elongation and strengthening is vastly limited. Here, large-scale glycome profiling coupled with fiber phenotype and transcriptome profiling was conducted on fiber collected daily covering the most critical fiber developmental window. High temporal resolution profiling allowed us to identify specific polysaccharide epitopes associated with distinct fiber phenotypes that might contribute to fiber quality. This study revealed the critical role of highly branched RG-I pectin epitopes such as, β-1,4-linked-galactans, β-1,6-linked-galactans, and arabinogalactans, in addition to earlier reported homogalacturonans and xyloglucans in the formation of cotton-fiber-middle-lamella and contributing to fiber plasticity and elongation. We also propose the essential role of heteroxylans (Xyl-MeGlcA and Xyl-3Ar), as a guiding factor for secondary CW cellulose-microfibril arrangement, thus contributing to fiber strength. Correlation analysis of glycome and transcriptome data identified several key putative glycosyltransferases involved in synthesizing the critical polysaccharide epitopes. Novel details discovered here provide a foundation to identify molecular factors that dictate important fiber traits.

## Introduction

Cotton (*Gossypium* species) is the most widely used natural textile fiber globally, with an estimated value of US $64 billion annually (USDA, 2024). Four cotton species have been domesticated, and among them *Gossypium hirsutum* is most widely planted because of its high yield and environmental adaptability (Hu et al. 2021; Jareczek et al. 2023; Viot and Wendel 2023). Cotton fiber trichomes reach about 2.5 to 4 cm in length during 45-50 days post-anthesis (DPA). The matured fiber is a dried cell wall (CW) containing about 95% cellulose and traces of non-cellulosic polysaccharides like callose, pectins, hemicelluloses, dried glycoproteins, sugars, and minerals (Haigler et al. 2012). During development, the CW polysaccharides are synthesized and dynamically undergo remodeling, which results in finer industrial-quality fibers. The quality of the fiber varies between cotton species, which is determined by fiber length and strength (Kim and Triplett 2001).

Fiber development is an intricately regulated process that is poorly understood. However, there are gross morphological landmarks that have been used to define key events. It is often divided into five sequential and overlapping events: initiation and tapering, elongation, transition, secondary CW (SCW) deposition, and desiccation/maturation (Haigler et al. 2012; Stiff and Haigler 2016). Unicellular trichoblasts (fiber initials) bud from the seed surface at -1 DPA, and undergo tapering and it is a precise cytoskeleton-dependent CW remodeling phase, which is completed by 2 DPA (Stiff and Haigler 2016; Graham and Haigler 2021; Yanagisawa et al. 2022). Primary CW (PCW) synthesis occurs over the first two-three weeks, during which cotton fiber develops its characteristic length and fineness (Meinert and Delmer 1977; Applequist et al. 2001). Fiber elongation rates begin to slow early in development, well before the transition to SCW synthesis (Schubert et al. 1973; Howell et al. 2024 unpublished). During the transition stage (16-20 DPA), cessation of PCW, beginning of SCW cellulose accumulation, and CW thickening occur. Drastic increase in SCW cellulose deposition, as well as CW thickening, happens during 16-35 DPA. Thereafter, the cytoplasm dries out, and the dried matured fibers twist, which makes them easily spun into threads (Kim and Triplett 2001; Haigler et al. 2012).

Cotton fiber development is a complex process, involving the coordination of gene expression networks, hormone signaling, physiology, and biosynthesis that determine CW polysaccharide composition, which in turn determines fiber quality (Haigler et al. 2012; Jan et al. 2022; Jareczek et al. 2023). Several studies were conducted on fiber CW polysaccharides which include, monosaccharide composition analysis, glycan microarrays-polysaccharide linkage analysis, microscopy techniques involving staining and immunolocalization, and glycome profiling with antibodies recognizing plant polysaccharide epitopes (Singh et al. 2009; Avci et al. 2013; Hernandez-Gomez et al. 2015; Hernandez-Gomez et al. 2017; Guo et al. 2019; Pettolino et al. 2022). These and other studies demonstrated that the fiber CW composition changes dynamically throughout development, especially during elongation and the transition to SCW synthesis. Nevertheless, these studies were conducted on a small-scale, and the information is limited specifically on the molecular dynamics of non-cellulosic polysaccharides during fiber development.

In young fiber between 0 and 2 DPA, a pectin-rich, xyloglucan-depleted outer sheath develops. By 4 DPA, the synthesis of a cotton fiber middle-lamella (CFML) is formed. The CFML serves to join adjacent fibers into tissue-like, highly organized, and tightly packed bundles, which eventually facilitates fiber elongation smoothly within the confined boll space (Haigler et al. 2012). Recent studies on glycome profiling of fiber with numerous monoclonal antibodies recognizing polysaccharide epitopes reported the presence of both fucosylated- and non-fucosylated xyloglucans, pectic arabinans/galactans, and homogalacturonans in CFML (Vaughn and Turley 1999; Singh et al. 2009; Avci et al. 2013; Hernandez-Gomez et al. 2017; Guo et al. 2019), and callose, cellulose, and heteromannans at PCW (Hernandez-Gomez et al. 2015).

During the growth phase, the impact of fiber turgor pressure, expansin (CW loosening protein), expression of a broader class of CW-synthesizing/remodeling enzymes, and the dynamics of polysaccharides in PCW and CFML, and cellulose microfibril decides the fate of cotton fiber plasticity, expansion, length, and spinnability, in different species (Shimizu et al. 1997; Orford and Timmis 1998; Harmer et al. 2002; An et al. 2007; Michailidis et al. 2009; Shao et al. 2011; Keynia et al. 2022; Yanagisawa et al. 2022).

During the transition stage, transcriptionally regulated CW degrading enzymes break down the CFML, which leads to reduction of pectin and xyloglucan molecular mass and separation of fibers (Meinert and Delmer 1977; Tokumoto et al. 2002; Singh et al. 2009; Guo et al. 2019). Also during transition stage, a unique “winding” CW layer is formed with differently oriented cellulose microfibrils than the microfibrils usually seen in PCW (Seagull, 1993). Also, during the transition stage, callose deposition reaches a peak, and it was proposed that both the callose and cellulose might be involved in the formation of the winding layer and contributing to fiber strength (Kerr 1946; Hsieh et al. 1995; Hinchliffe et al. 2011).

At the onset of SCW thickening, the rate of cellulose synthesis increases drastically, and both microtubules and cellulose fibrils adopt a steep helix relative to the fiber axis (Meinert and Delmer 1977; Maltby et al. 1979; Seagull, 1993; Martin and Haigler 2004; Abidi et al. 2010). The SCW mainly contains cellulose, and its content and biophysical properties, such as degree of crystallinity, fibril angle-twist, and degree of polymerization mainly decide the strength of the fiber (Haigler et al. 2012).

Earlier research clearly points out that the fiber elongation and the transition stage (interplay of elongation and the beginning of SCW thickening), are the most crucial stages that determine the outcome of fiber quality (Haigler et al. 2012; Tuttle et al. 2015; Jan et al. 2022; Jareczek et al. 2023). So far, only a few small-scale studies have been conducted on non-cellulosic polysaccharides with few DPA time points, and rarely included transcriptome analysis. In the present study, large-scale glycome and transcriptome profiling was carried out in parallel along with fiber-phenotyping on fibers purified daily from 6 to 25 DPA. A broad collection of carbohydrate-specific monoclonal antibodies was utilized in combination with monosaccharide analyses. This study has discovered CW polysaccharide epitopes that could be key factors contributing to fiber length and strength. Correlation analysis of glycomic and transcriptomic data identified target genes that are predicted to influence cotton fiber CW composition.

## Results

### Polysaccharide and monosaccharide composition during cotton fiber development (6 - 25 DPA)

Alcohol insoluble CW, dried-lyophilized buffer soluble pectin-enriched-fraction (50mM CDTA:50mM ammonium oxalate buffer extracts), alkali-soluble hemicellulose-enriched-fraction (4M KOH extracts), and total cellulose (amorphous and crystalline) from 6 to 25 DPA cotton fibers were each weighed and summarized (Fig. 1A; Supplemental Fig. S1; Supplemental Table S1). As expected, the weight of total CW content and content of polysaccharide fractions per boll increased gradually from 6 to 25 DPA (Supplemental Table S1A). However, percentage wise, the pectin, and hemicellulose, in the total CW slowly started decreasing from 32.7%, and 22.3% at 6 DPA and reached around 6.6% and 7.3% by 25 DPA, respectively (Fig. 1A; Supplemental Table S1B). In contrast, the total cellulose increased gradually from 26.4% at 6 DPA, reached 40% at 16 DPA and increased drastically afterwards to reach 81.9% at 25 DPA. The crystalline cellulose content was 2.6% at 6 DPA and reached 13.6% at 16 DPA. After 16 DPA, it rapidly increased, and reached 52.4% by 25 DPA.

**Figure 1.**
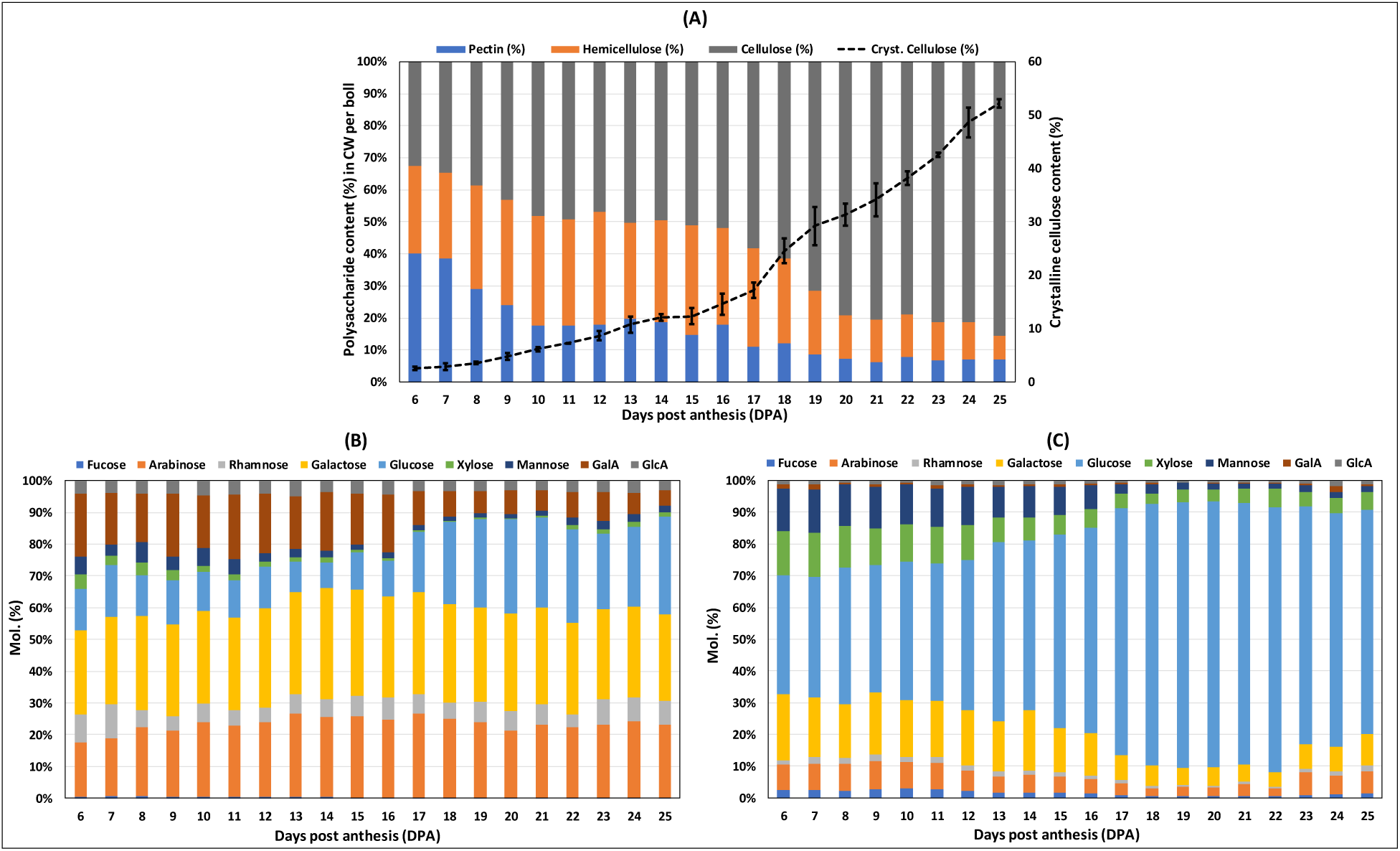
Polysaccharide content and their monosaccharide composition of cotton fiber cell walls during development (6 - 25 DPA). **A)** Polysaccharide content of cotton fibers cell wall (CW). Cotton fiber CW content was fractionated into pectin, hemicellulose and cellulose polysaccharides, dried and weighed. The graph represents the polysaccharides content in % and it is calculated by keeping the CW weight as 100 %. **B)** Monosaccharides content of the pectin-enriched fractions (in Mol%). **C)** Monosaccharides content of the hemicellulose-enriched fractions (in Mol%). The data are average from three biological replicates.

Monosaccharide composition of pectin-enriched-fractions showed that the relative content of arabinose slightly increased (17% to 23%) and glucose significantly increased (13% to 31%) between 6 and 25 DPA (Fig. 1B; Supplemental Table S2A). The content of the xylose (4.6% to 1.6%), mannose (5.4% to 2.3%), fucose (0.6% to 0.2%), and GalA (20% to 5.1%) significantly decreased between 6 and 25 DPA. The monosaccharide composition of hemicellulose-enriched-fractions showed that glucose content (38.5% to 74.2%) significantly increased from 6 to 25 DPA (Fig. 1C; Supplemental Table S2B). The content of arabinose slightly decreased (7.1% to 2.6%) between 6 and 22 DPA. Galactose (19.4% to 8.5%), xylose (13.9% to 4.7%), and mannose (14.4% to 1.5%) showed significant reductions between 6 and 25 DPA.

### Glycome profiling and self-organizing maps (SOM) of diverse epitope patterns of the polysaccharides

Seventy-one different antibodies were used to profile cotton fiber polysaccharide epitopes (Fig. 2; Supplemental Table S3). Since the ELISA data for each epitope originate from two different fractions of the same sample, for practicality, the naming of epitopes was suffixed with either “- P” for pectin-enriched-fractions or “-HC” for hemicellulose-enriched-fractions. The glycome data provided the temporal variability of CW composition as a function of developmental time. A challenge in this form of data analysis is to objectively define groups of epitopes that share similar temporal profiles. Using the data obtained from the glycome profiling of pectin-enriched- and hemicellulose-enriched-fractions (Supplemental Tables S4 and S5), SOM-based clusters and heat maps were generated to visualize and classify the polysaccharide epitope distribution patterns (Fig. 3; Supplemental Table S6). SOM for the pectin-enriched-fractions showed that most of the polysaccharide epitope patterns started with a higher amount at around 6 DPA, and gradually reduced until 25 DPA (Fig. 3A). In contrast, the SOM for the hemicellulose-enriched-fractions showed more variability in the epitope patterns (Fig. 3C). The heat maps displayed clearly that the epitope profile pattern was reproducible among all developmental stages of all the three biological replications sampled. Both the SOM and heat maps conclusively allowed us to group the epitopes further into four “categories” (Tables 1 to 4; Figs. 4 to 7).

**Figure 2.**
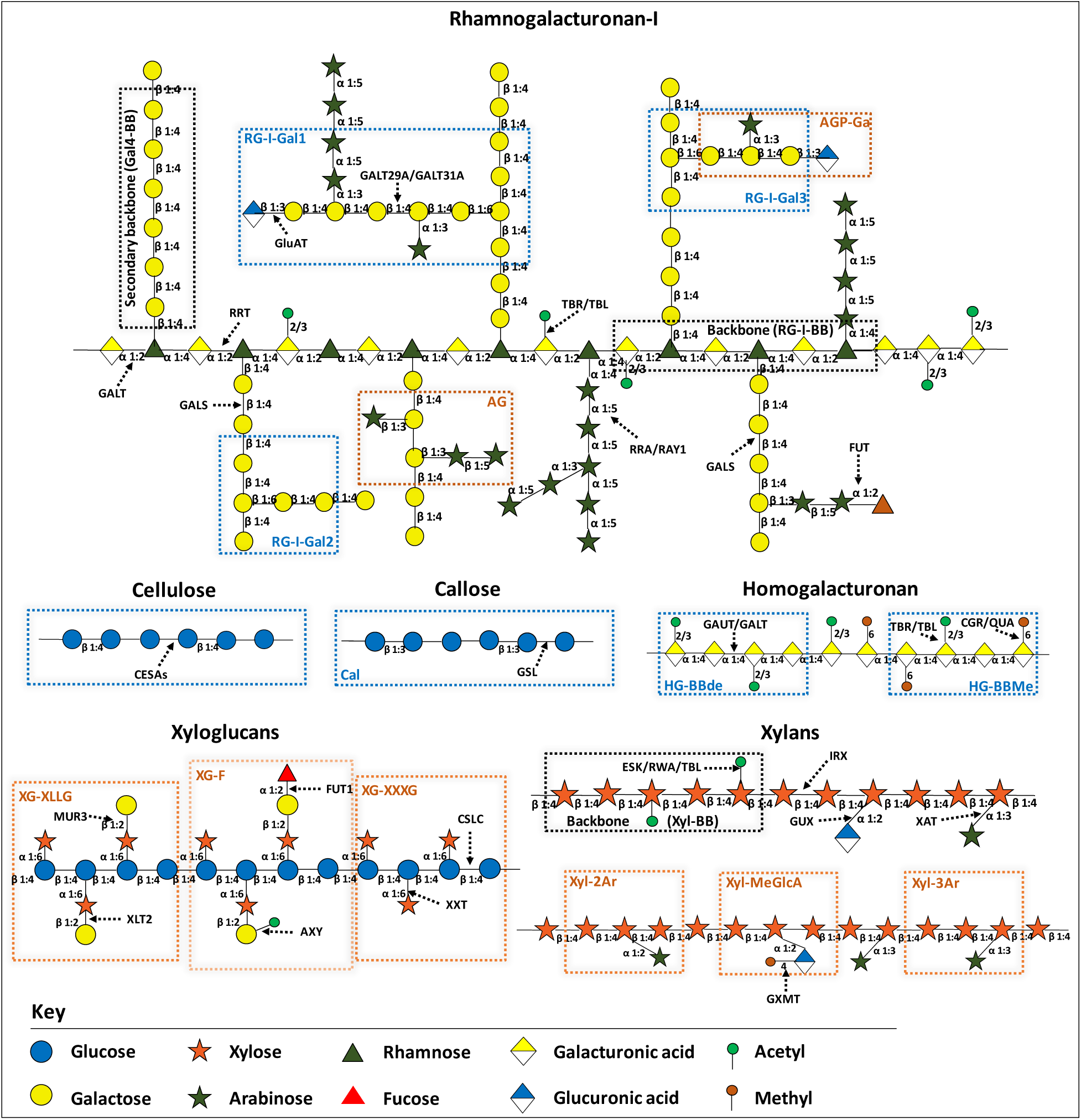
Pictorial representation of structures of analyzed cell wall (CW) polysaccharides epitopes and glycosyltransferase enzymes involved in their synthesis. The schematic structure of Rhamnogalacturonan-I (RG-I) pectic polysaccharide displays different epitopes like, backbones (RG-I-BB, Gal4-BB), β-1,6-linked galactans (RG-I-Gal1, RG-I-Gal2 and RG-I-Gal3), and arabinogalactans (AG, AGP-Ga). Cellulose and callose polysaccharides are made up of chain of glucose molecules connected by β-1,4- and β-1,3-linkages, respectively. Homogalacturonan (HG) pectic polysaccharide displays two different epitopes, Methyl-esterified HG (HG-BBMe), and de-esterified HG (HG-BBde). Xyloglucan (XG) structure shows, xylosylated XG (XG-XXXG), galactosylated XG (XG-XLLG), and fucosylated XG (XG-F) epitopes. The schematic structures of xylans show the backbone (Xyl-BB), arabinoxylans (Xyl-2Ar and Xyl-3Ar) and methylated-glucuronoxyaln (Xyl-MeGlcA) epitopes. Refer to Supplemental Table S3 for the details of the epitopes recognized by the antibodies used in this study. The known or putative Arabidopsis CW synthesizing enzymes are denoted by black dotted arrows.

**Table 1.**
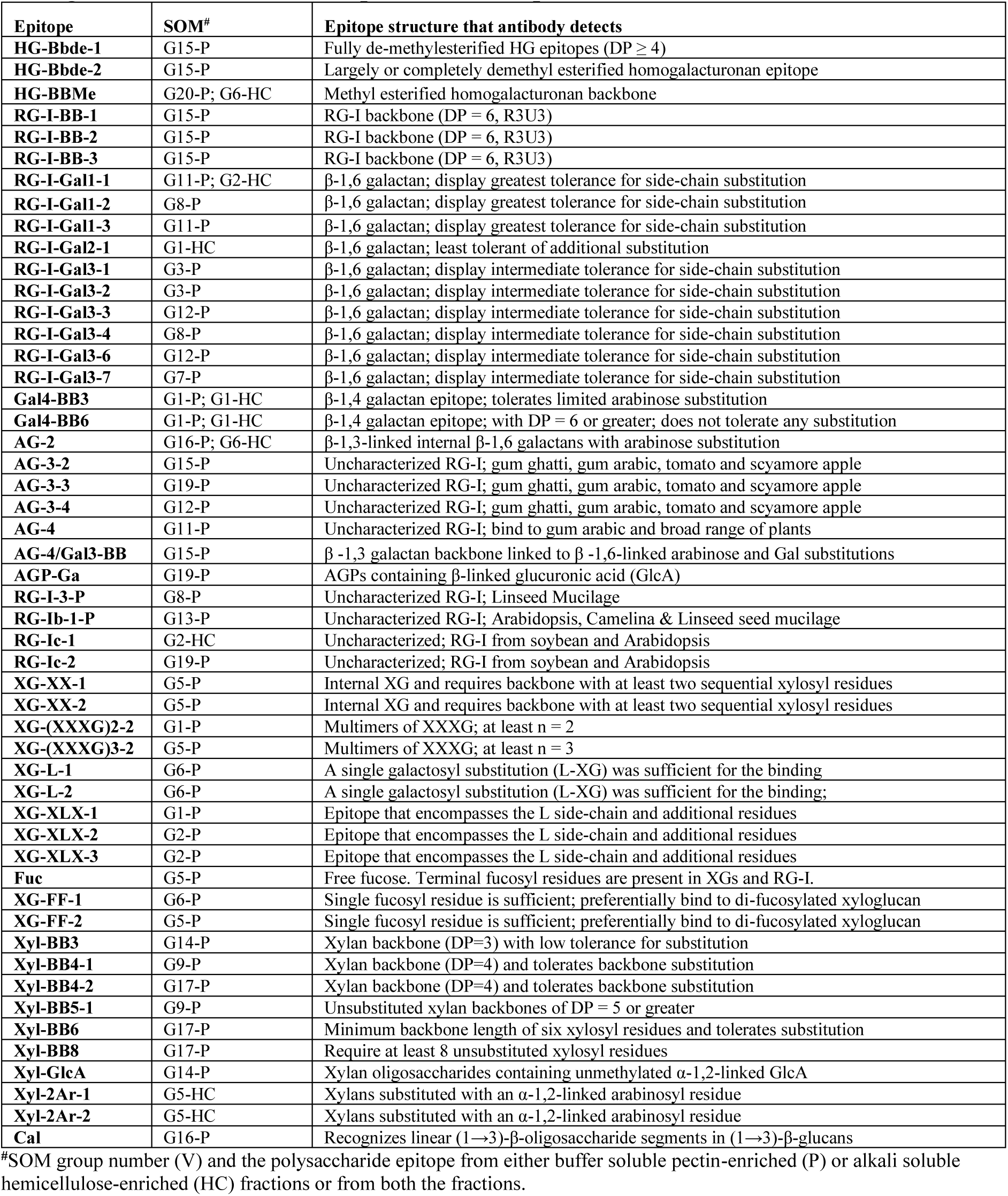
List of polysaccharide epitopes that are highly abundant at early stages and reduced during *G. hirsutum* fiber development (6 - 25 dpa).

The first category includes 51 different polysaccharide epitopes that are highly abundant at early stages and reduced with varying slopes and timings to low levels by 25 DPA (Table 1; Fig. 4). Most of the buffer-soluble pectin, xyloglucan, and xylan epitopes displayed this pattern. Interestingly, each of these three polysaccharides distinctly formed three different separate subgroups within the first category. For example, most of the pectin epitopes located in a set of nearby SOM groups and formed a distinct subgroup (SOM groups G3-P, G7-P, G8-P, G11-P, G12-P, G15-P, G16-P, G19-P; Figs. 3A and 4A; Supplemental Table S6A), where their content declined slowly starting from 12 DPA. On the contrary, the content of most of the xyloglucan (SOM groups G1-P, G2-P, G5-P, G6-P; Figs. 3A and 4B) and RG-I-β-1,4-linked-galactan epitopes (SOM group G1-P; Figs. 3A and 4B) reduced drastically around 16 DPA, coinciding with the transition stage to SCW synthesis, which forms the second subgroup. The third subgroup mainly has the xylan epitopes (SOM groups G9-P, G10-P, G13-P, G14-P, G17-P; Figs. 3A and 4C), and their content reduced slowly and gradually from the beginning of 6 DPA.

**Figure 3.**
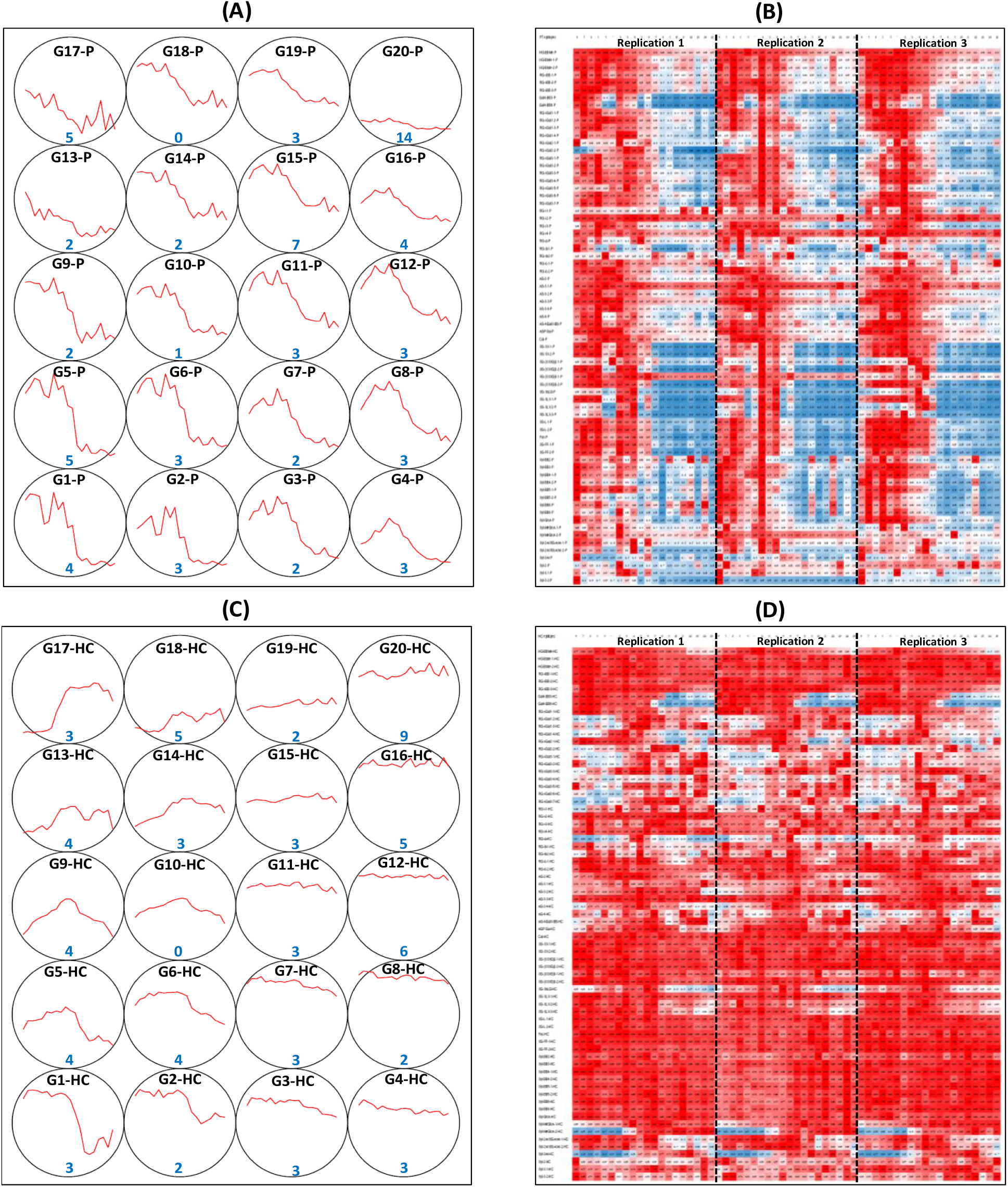
Self-organizing map (SOM) and heat map of the epitope patterns obtained from glycome profiling. **A, B)** Computer generated SOM of the glycome epitope patterns of buffer soluble pectin-enriched (50mM CDTA-50mM Ammonium oxalate extract) polysaccharide fractions and the corresponding heat map, respectively. **C, D)** SOM of alkali soluble hemicellulose-enriched (4M KOH extract) polysaccharide fractions and the corresponding heat map, respectively. SOM shows that there are 20 groups (G1 to G20) within each fraction and the number in blue font denotes the number of epitopes falls within each SOM group. The data are average from three biological replicates. Epitopes from either buffer-soluble pectin-enriched (50mM CDTA-50mM Ammonium oxalate extract) or alkali-soluble hemicellulose-enriched (4M KOH extract) fractions are denoted by the suffix “-P” and “-HC”, respectively. Refer to Supplemental Table S6 for the details of epitopes falls within each SOM group. Heat maps were generated on excel and the epitope abundances were standardized from 0 (blue) to 1 (red). Heat maps show the consistency of epitopes content from 6 to 25 DPA between the three replicates sampled.

**Figure 4.**
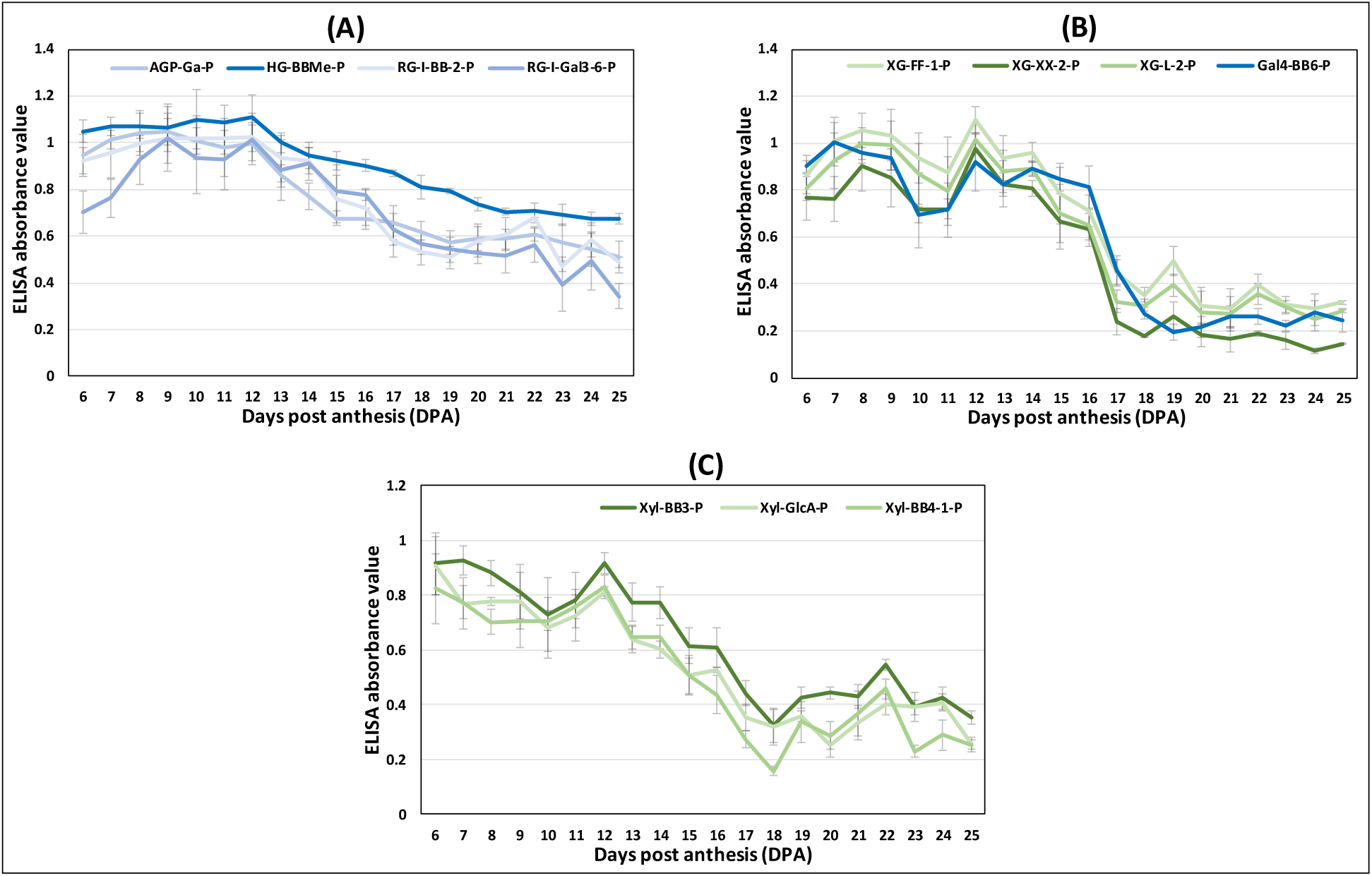
Glycome profile of polysaccharide epitopes that are highly abundant at early stages and reduced during *G. hirsutum* fiber development (6 - 25 DPA). **A)** Profile of representative pectin epitopes listed in Table 1 that decreased slowly. **B)** Profile of representative xyloglucan (XG) or pectin epitopes listed in Table 1 that decreased drastically at 16 DPA. **C)** Profile of representative xylan (xyl) epitopes listed in Table 1 that decreased at faster rate gradually. The data are mean from three biological replications along with standard error bars. Epitopes from either buffer soluble pectin-enriched (50mM CDTA-50mM Ammonium oxalate extract) or alkali soluble hemicellulose-enriched (4M KOH extract) fractions are denoted by the suffix “-P” and “-HC”, respectively. The pectin and hemicellulose epitopes profile are color coded by blue and green, respectively.

Another set of SOM group containing fourteen of the epitopes (all from the hemicellulose-enriched fraction) were placed in a second category (Table 2; Fig. 5). The content of these epitopes was either low at early DPAs and then rapidly increased around 12 DPA (Fig. 5A) or gradually increased from the beginning (Fig. 5B) to a higher level at later stages. Highly branched RG-I epitopes (β-1,6 linked galactans and arabinogalactans; SOM groups G13-HC, G14-HC, G17-HC, G18-HC) and three of the xylan epitopes (SOM groups G14-HC, G17-HC) constitute this second category (Figs. 3C and 5; Supplemental Table S6B).

**Table 2.** List of polysaccharide epitopes that are low at early stages and increased during *G. hirsutum* fiber development (6 - 25 dpa).

| Epitope | SOM# | Epitope structure that antibody detects |
| --- | --- | --- |
| **RG-I-Gal1-2** | G13-HC | β-1,6 galactan; display greatest tolerance for side-chain substitution |
| **RG-I-Gal1-3** | G13-HC | β-1,6 galactan; display greatest tolerance for side-chain substitution |
| **RG-I-Gal2-2** | G14-HC | β-1,6 galactan; least tolerant of additional substitution |
| **RG-I-Gal3-1** | G18-HC | β-1,6 galactan; display intermediate tolerance for side-chain substitution |
| **RG-I-Gal3-3** | G14-HC | β-1,6 galactan; display intermediate tolerance for side-chain substitution |
| **RG-I-Gal3-6** | G18-HC | β-1,6 galactan; display intermediate tolerance for side-chain substitution |
| **RG-I-Gal3-7** | G17-HC | β-1,6 galactan; display intermediate tolerance for side-chain substitution |
| **AG-3-2** | G9-HC | Uncharacterized RG-I; gum ghatti, gum arabic, tomato and scyamore apple |
| **AG-4** | G13-HC | Uncharacterized RG-I; bind to gum arabic and broad range of plants |
| **AG-4/Gal3-BB** | G13-HC | β-1,3 galactan backbone linked to β-1,6-linked arabinose and Gal substitutions |
| **Xyl-MeGlcA-1** | G14-HC | Xylans containing O-4 methylated α-1,2-linked GlcA |
| **Xyl-MeGlcA-2** | G17-HC | Xylans containing O-4 methylated α-1,2-linked GlcA |
| **Xyl-3Ar** | G17-HC | ExclusiGe to xylan consisting of single α-1,3-linked arabinofuranosyl residue |

#SOM group number (G) and the polysaccharide epitope from alkali soluble hemicellulose-enriched (HC) fractions.

**Figure 5.**
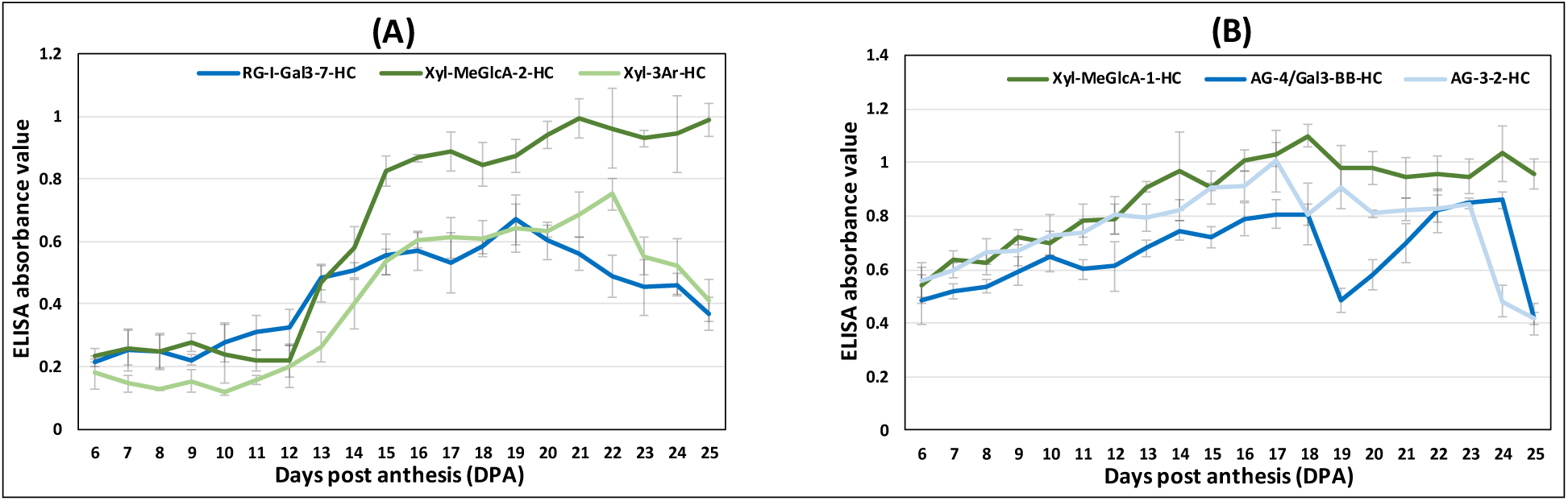
Glycome profile of polysaccharide epitopes that are low at early stages and increased during *G. hirsutum* fiber development (6 - 25 DPA). **A)** Profile of representative epitopes listed in Table 2 that increased drastically at 12 DPA. **B)** Profile of the epitopes listed in Table 2 that increased gradually from the beginning. The data are mean from three biological replications along with standard error bars. Epitopes from alkali-soluble hemicellulose-enriched (4M KOH extract) fractions are denoted by the suffix “-HC”. The pectin and hemicellulose epitopes profile are color coded by blue and green, respectively.

Some of the epitopes showed bell or semi-bell-shaped patterns, and their content reached a peak at mid-points (either 12 or 16 DPA) in the sampling window. Seven of such epitopes were grouped in the third category (Table 3; Fig. 6). The content of three epitopes, all came from a pectin-enriched fraction (SOM groups G4-P, G7-P; Figs. 3A and 6A), peaked at 12 DPA. Similarly, the content of the other four epitopes from a hemicellulose-enriched fraction (SOM groups G5-HC, G9-HC; Figs. 3B and 6B) peaked at 16 DPA. The fourth category of polysaccharide profiles includes 30 epitopes (7 pectins, 13 xyloglucans, 9 xylans, and 1 callose; all from the hemicellulose-enriched-fraction), which are present in a set of nearby SOM groups (SOM group G3-HC, G4-HC, G7-HC, G8-HC, G11-HC, G12-HC, G15-HC, G16-HC, G20-HC; Table 4; Figs. 3C and 7). These epitopes showed high abundance, and their patterns statistically remain unchanged between 6 and 25 DPA.

**Table 3.** List of polysaccharide epitopes that are peaked high at mid-stages of *G. hirsutum* fiber deGelopment (6 - 25 dpa).

| Epitope | SOM<sup>#</sup> | Epitope structure that antibody detects |
| --- | --- | --- |
| **RG-I-Gal1-4** | G7-P; G5-HC | β-1,6 galactan; display greatest tolerance for side-chain substitution |
| **RG-I-Gal2-2** | G4-P | β-1,6 galactan; least tolerant of additional substitution |
| **RG-I-Gal3-5** | G4-P | β-1,6 galactan; display intermediate tolerance for side-chain substitution |
| **AG-3-4** | G9-HC | Uncharacterized RG-I; gum ghatti, gum arabic, tomato and scyamore apple |
| **AGP-Ga** | G9-HC | AGPs containing β-linked glucuronic acid (GlcA) |
| **RG-Ia** | G9-HC | Uncharacterized; selectiGe to gum karaya plant preparations |
| **XG-XnLG** | G5-HC | Non-reducing terminal X side-chain and the L side-chain closest to the reducing terminus |

<sup>#</sup>SOM group number (G) and the polysaccharide epitope from either buffer soluble pectin-enriched (P) or alkali soluble hemicellulose-enriched (HC) fractions or from both the fractions.

**Table 4.**
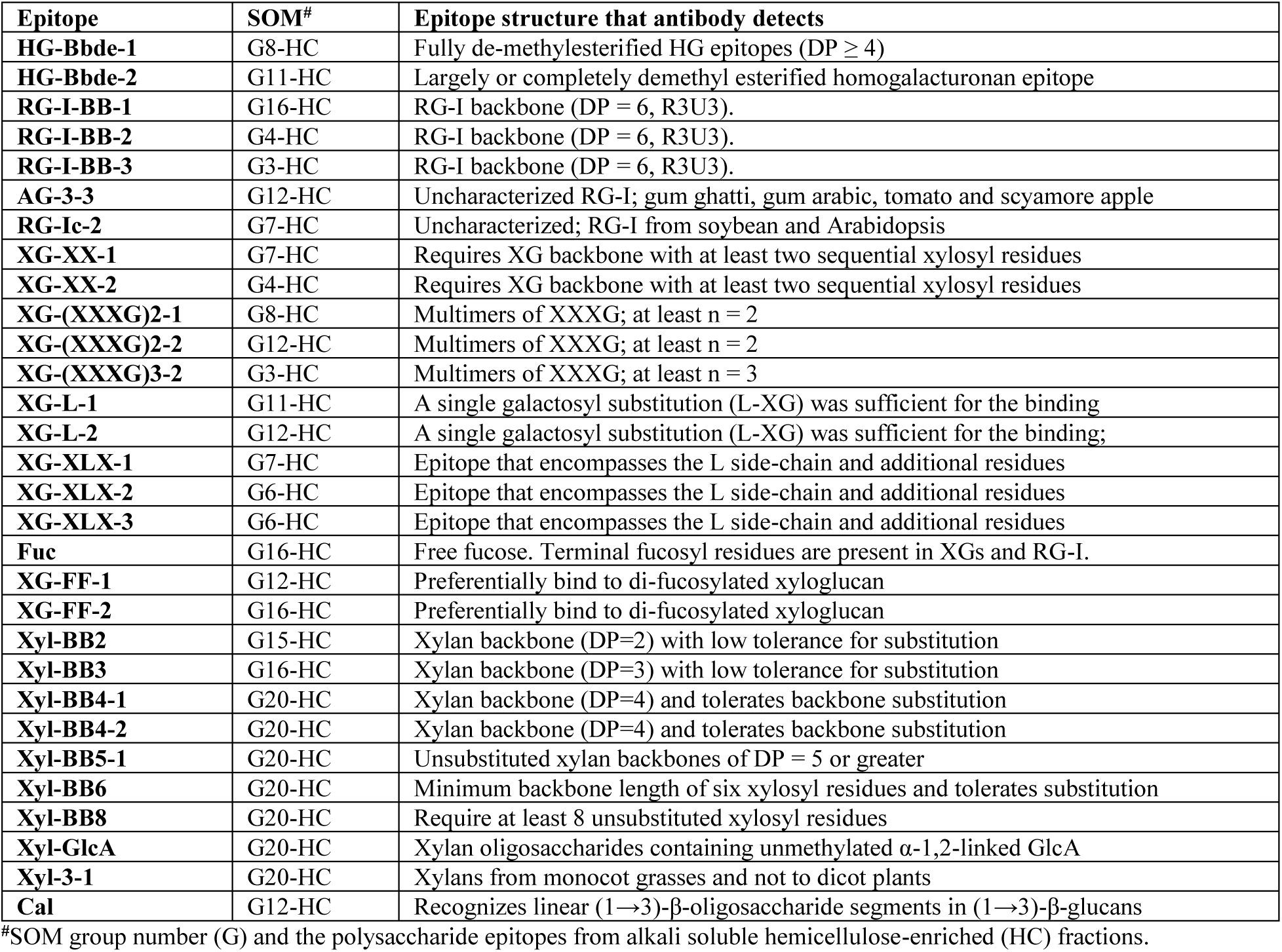
List of polysaccharide epitopes that are highly abundant and haGing horizontal pattern during *G. hirsutum* fiber deGelopment (6 - 25 dpa).

**Figure 6.**
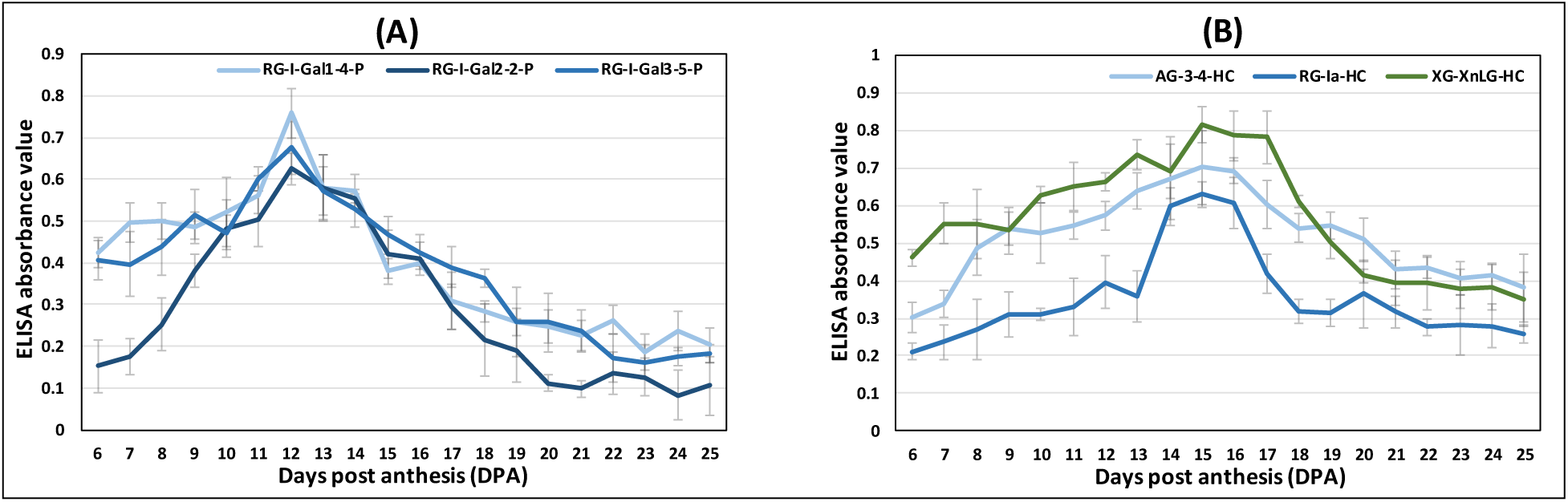
Glycome profile of polysaccharide epitopes that peaked high at mid-stage of *G. hirsutum* fiber development (6 - 25 DPA). **A, B)** Profile of polysaccharide epitopes listed in Table 3 reached peak at 12 DPA and 16 DPA, respectively. The epitopes content was low at the earlier and later DPAs but peaked high at the mid-stage, resembling a bell or semi-bell-shaped curve. The data are mean from three biological replications along with standard error bars. Epitopes from either buffer-soluble pectin-enriched or alkali-soluble hemicellulose-enriched fractions are denoted by the suffix “-P” and “-HC”, respectively. The pectin and hemicellulose epitopes profile are color coded by blue and green, respectively.

**Figure 7.**
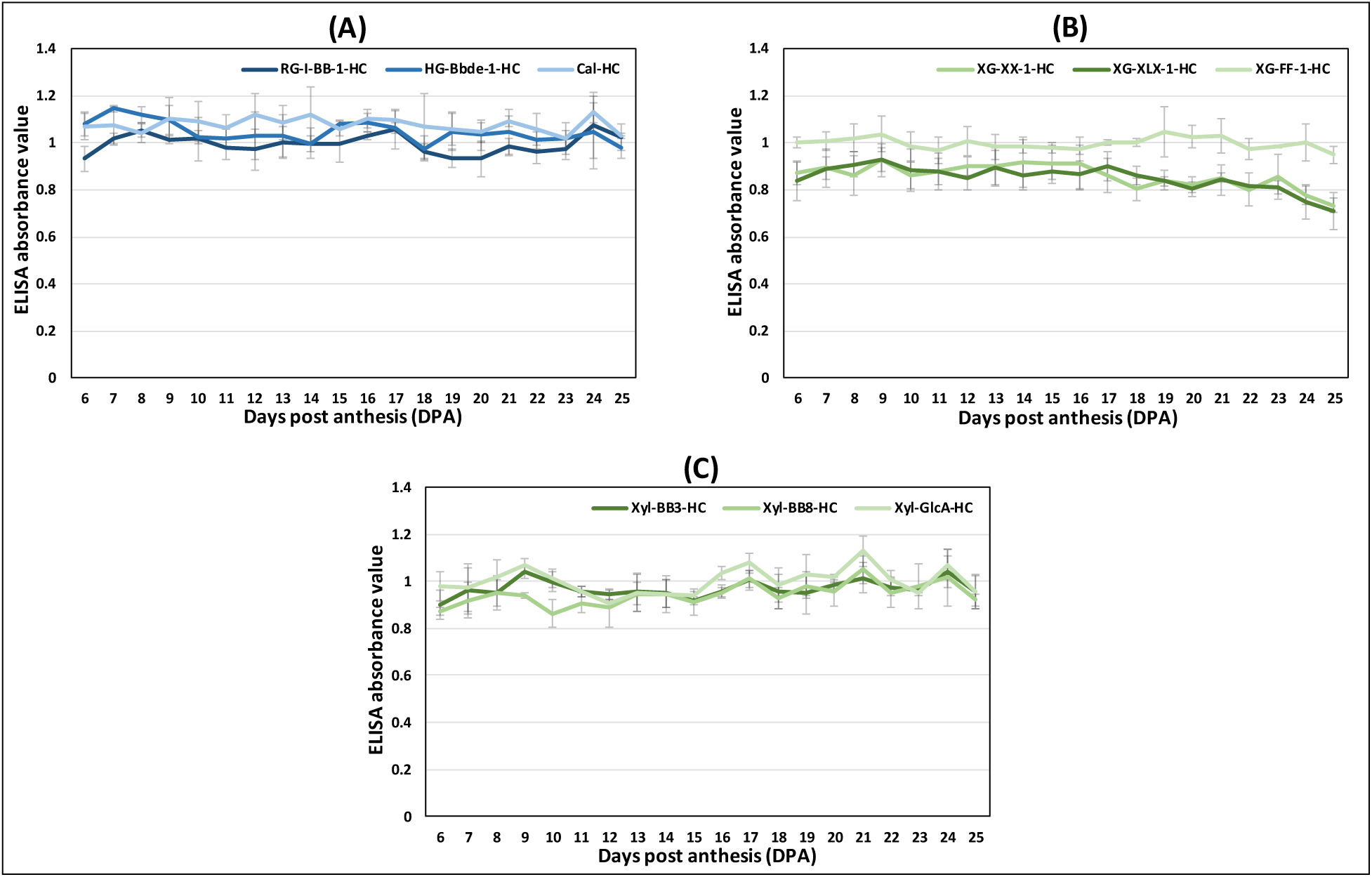
Glycome profile of polysaccharide epitopes having high abundance and horizontal pattern during *G. hirsutum* fiber development (6 - 25 DPA). **A, B, C**) Profile of the representative pectin (HG/RG-I)/callose (Cal), xyloglucan (XG) and xylan (Xyl) polysaccharide epitopes listed in Table 4, respectively. The epitopes content was high throughout the 6 - 25 DPA. The data are mean from three biological replications along with standard error bars. Epitopes from alkali-soluble hemicellulose-enriched fractions are denoted by the suffix “-HC”. The pectin and hemicellulose epitopes profile are color coded by blue and green, respectively.

### Correlation analysis of cotton fiber polysaccharide epitopes and fiber phenotypes

Pearson correlation coefficient (PCC) analyses were carried out between the profiles of glycome epitopes and profiles of fiber phenotypes generated simultaneously from a related collaborative study (Howell et al. 2024 unpublished) to predict the potential epitopes involved in governing the phenotypes. Phenotypes such as fiber elongation rate, turgor pressure, microfibril orientation, and cellulose content were used, and a PCC cut-off value of ≥ 0.7 was considered to be a strong positive correlation (Supplemental Table S7; Supplemental Fig. S2A).

PCC results revealed that many of the pectin and buffer-soluble xyloglucan and xylan epitopes categorized in Table 1 significantly correlated and closely mirrored the decreasing fiber growth rates from 6 to 25 DPA (Fig. 8A; Supplemental Table S7). Few of the pectin and xyloglucan epitopes highly correlate with the fiber turgor pressure profile, which peaked at 16 DPA and reduced drastically to a low level at later stages (Fig. 8B; Supplemental Table S7).

**Figure 8.**
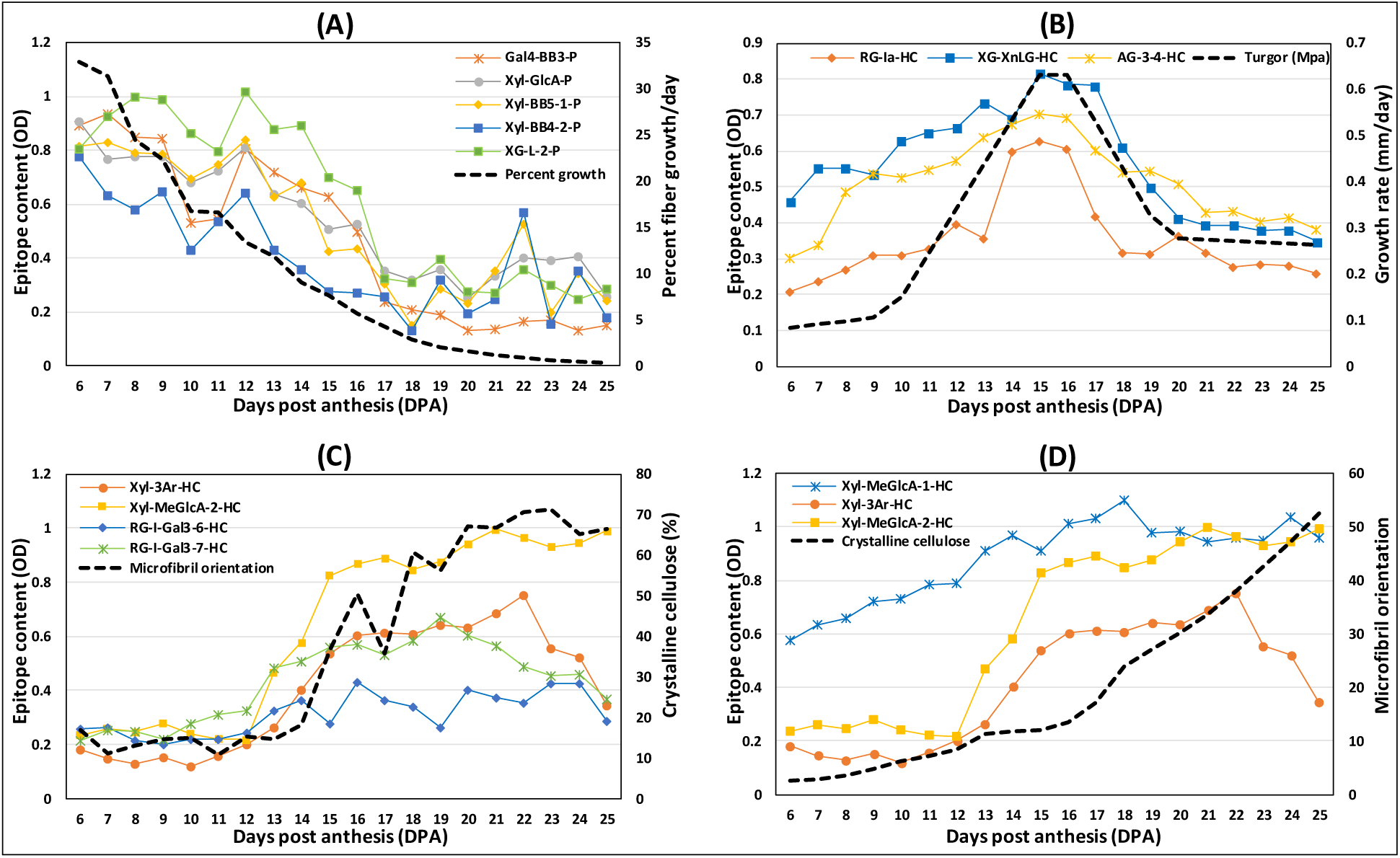
Profiles of fiber phenotypes and the correlated polysaccharide epitopes. **A)** Profile of percentage fiber growth rate and few of the representative correlated polysaccharide epitopes. **B)** Profie of fiber tugor pressure and few of the representative correlated polysaccharide epitopes. **C)** Profile of microfibril orientation and the correlated polysaccharide epitopes. **D)** Profile of fiber cell wall crystalline cellulose and the correlated polysaccharide epitopes. Refer to Supplemental Table S7 for the full list of correlated and non-correlated epitopes.

The profile of microfibril orientation rapidly changes around 16 DPA, coinciding with transition to SCW synthesis (Fig. 8C; Howell et al. 2024 unpublished). Profiles of three of the xylan epitopes (Xyl-MeGlcA-1-HC, Xyl-MeGlcA-2-HC and Xyl-3Ar-HC) and two of the highly branched RG-I epitopes (RG-I-Gal3-6-HC and RG-I-Gal3-7-HC) were highly correlated with the profile of microfibril orientation (Fig. 8C; Supplemental Table S7). Interestingly, the profile of microfibril orientation is similar to the profile of cellulose content (Fig. 8D). The cellulose content drastically increased after 16 DPA coinciding with the beginning of SCW synthesis, and here also the profiles of three of the above mentioned xylan epitopes highly correlated with the cellulose content profile (Fig. 8, C and D; Supplemental Table S7).

### Correlation analysis of cotton fiber polysaccharide epitopes and transcripts of enzymes potentially involved in the biosynthesis of the epitopes

*Gossypium hirsutum* is an allotetraploid, and its gene families tend to be large. Therefore, it is challenging to predict which particular gene/protein might be responsible for the observed developmental profile of a CW epitope. However, to predict the most likely gene(s), correlation analyses were conducted using the glycome data and transcriptome data collected at the same time points. Observing a high correlation between transcript expression pattern and polysaccharide epitope abundance would suggest the potential involvement of particular enzymes in the synthesis of specific epitopes.

First, the list of *Arabidopsis* enzymes known or predicted to be involved in the biosynthesis of CW polysaccharides was generated (Supplemental Table S8A). Then, using *Arabidopsis* gene sequences, corresponding homologous *G. raimondii* genes were identified from the Phytozome (www. https://phytozome-next.jgi.doe.gov/). Using this information, corresponding genes were pulled out from the transcriptome data generated in our study (Supplemental Table S8B). Next, the PCC analyses were carried out (Supplemental Fig. S2B).

#### i) Correlation analysis of homogalacturonan (HG) related epitopes and the corresponding transcripts

In *Arabidopsis* and some other plants, 15 galacturonosyltransferases (GAUTs) and 10 galacturonosyltransferase-like (GALTs) enzymes are involved in HG backbone biosynthesis by adding galacturonic acid (GalA) to the growing chain of HG backbone. There are also HG methyl transferases (CGRs/QUAs) and O-acetyltransferases (TBLs/TBRs) involved in decorating the HG backbone with methyl and acetyl groups, respectively (Atmodjo et al. 2013; Engle et al. 2022) (Fig. 2).

Our search for *Arabidopsis* homologous genes revealed that there are 48 GAUTs, 24 GALTs, 24 methyltransferases, and 14 acetyltransferases expressed in *G. hirsutum* fibers, which includes both A and D homoeologs (Supplemental Table S8B). The profiles of three of the HG epitopes (HG-BBde-1-P, HG-BBde-2-P, and HG-BBMe-P) were correlated with subsets of gene expression profiles from the genes in the above three classes of enzymes. The PCC correlations ranged from -0.9 to +0.9 in the entire gene set; however, the levels of 10 GAUTs, 3 GALTs, 6 methyltransferases, and 3 acetyltransferases had PCC values exceeded 0.7 cut-off and were defined as candidate genes involved in the observed HG profiles (Fig. 9A; Supplemental Table S8C).

**Figure 9.**
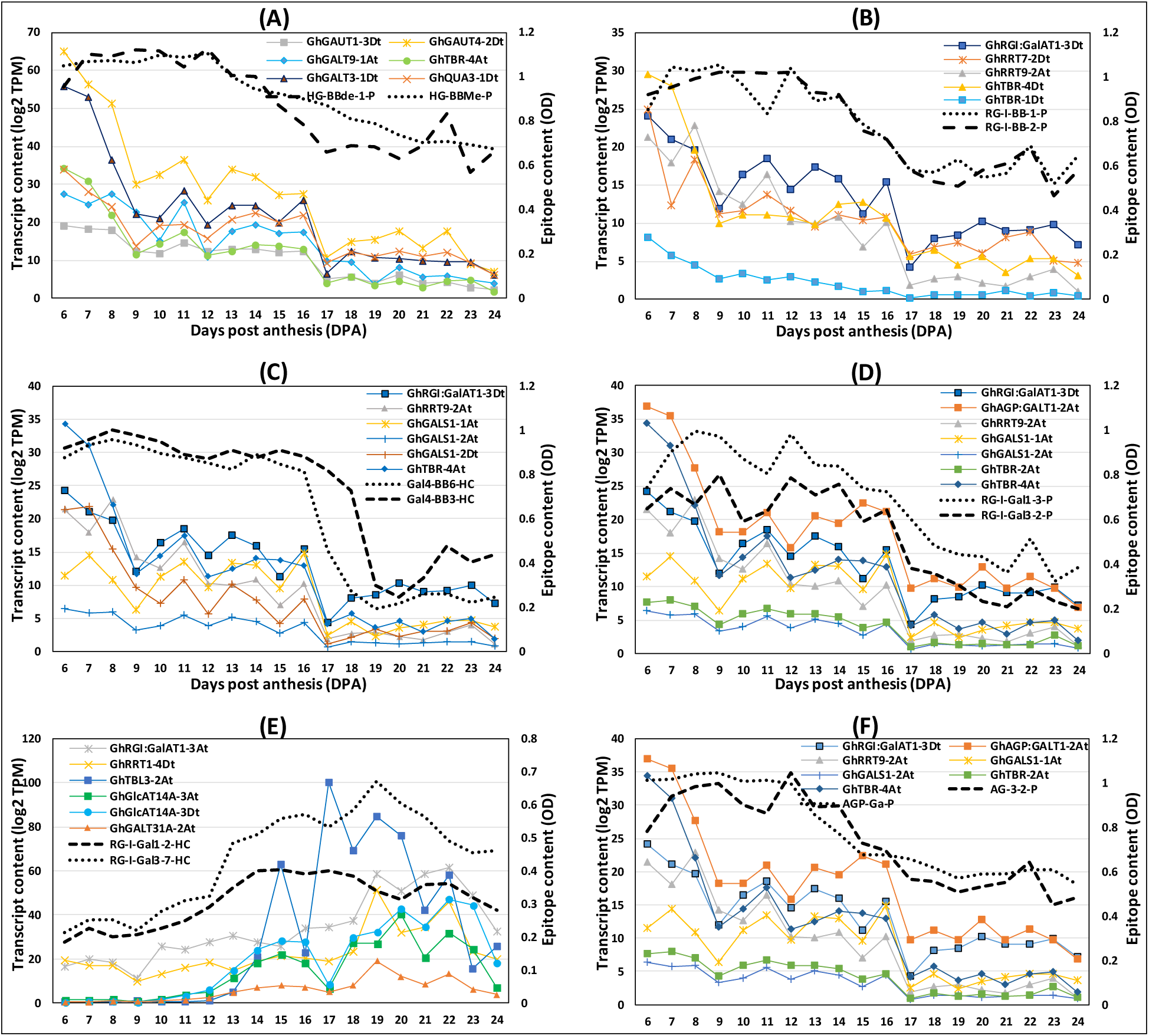
Profiles of pectin epitopes and the correlated profiles of transcripts of enzymes involved in synthesizing of these epitopes. **A)** Profile of homogalacturonan epitopes (HG-Bbde-P and HG-BBMe-P) and correlated representative transcripts. **B)** Profile of RG-I primary backbone epitopes (RG-I-BB1-P and RG-I-BB2-P) and correlated representative transcripts. **C)** Profile of RG-I secondary backbone β-1,4 linked galactan epitopes (Gal4-BB3-HC and Gal4-BB6-HC) and correlated representative transcripts. **D)** Profile of RG-I-β-1,6 linked galactan epitopes (content going down) (RG-I-Gal1-3-P and RG-I-Gal3-2-P) and correlated representative transcripts. **E)** Profile of RG-I-β-1,6 linked galactan epitopes (content going up) (RG-I-Gal1-2-HC and RG-I-Gal3-7-HC) and correlated representative transcripts. **F)** Profile of Arabinogalactan epitopes (AG-3-2-P and AGP-Ga-P) and correlated representative transcripts. Refer to Supplemental Table S8 (8B to 8H) for the full list of correlated and non-correlated transcripts. Refer to Fig. 2 for the details of epitope structures and the CW synthesizing enzymes involved.

#### ii) Correlation analysis of Rhamnogalacturonan-I (RG-I) related epitopes and the corresponding transcripts

The RG-I primary backbone is branched with the β-1,4-galactans, which constitute the secondary backbone of RG-I molecules. Further, the β-1,4-galactans are decorated with β-1,6-linked galactan, and arabinan side chains (Fig. 2). Galacturonosyltransferases (RG-I:GalATs), RG-I:rhamnosyltransferases (RRTs), and O-acetyltransferases (TBR/TBL) are required to synthesize the primary backbone of RG-I (Fig. 2) (Atmodjo et al. 2013; Amos et al. 2022). β-1,4-galactan synthases (GALSs) synthesizes β-1,4-galactan secondary backbone of RG-I. The AGP-GALTs synthesize the β-1,3-galactan secondary backbone of RG-I or arabinogalactan protein (AGP). RG-I and AGPs of CW share many structural epitopes, and it is hard to distinguish the origin of these epitopes from these two sources (Thorne et al. 2023). The β-1,6-galactosyltransferases (GALT29A/GALT31A) are involved in adding β-1,6-linked galactans to the secondary backbone. β-arabinofuranosyl transferases (RAY1) are known to participate in adding arabinose to β-1,6-linked galactans (Atmodjo et al. 2013).

The search for the cotton orthologs to *Arabidopsis* revealed that there are 8 RG-I:GalATs, 34 RRTs, 14 TBR/TBLs, 6 GALS, 34 AGP:GALTs, 10 GALT29A/GALT31A, and 2 RAY1 expressed in cotton fibers, including both A and D homoeologs (Supplemental Table S8B). Three of the RG-I primary backbone (RG-I-BB) epitopes profiles from the pectin-enriched-fraction were PCC analyzed with the transcript profiles of 8 RG-I:GalATs, 34 RRTs, and 14 TBR/TBLs, which are the enzymes involved in backbone synthesis. PCC analysis results showed that one RG-I:GalATs, three RRTs, and four TBRs highly correlated with the RG-I-BB epitopes (Fig. 9B; Supplemental Table S8D). In our study, two of the antibodies recognized the β-1,4-galactan secondary backbone epitopes (Gal4-BB3-HC and Gal4-BB6-HC). Both the epitopes and the transcript profile of the 8 RG-I:GalATs, 34 RRTs, 14 TBR/TBLs, and 6 GALS genes were PCC analyzed and it showed that 1 RG-I:GalAT, 4 RRTs, 3 GALSs, and 3 TBRs highly correlated with both the epitopes from β-1,4-galactan secondary backbone (Fig. 9C; Supplemental Table S8E).

In the present study, 13 antibodies were used to detect the RG-I-β-1,6-linked galactans which are linked to the β-1,3-galactan or β-1,4-galactan secondary backbones of RG-I. PCC analysis was conducted between 7 RG-I β-1,6-linked galactan epitopes (content goes down during 6-25 DPA) and the transcripts of 8 RG-I:GalATs, 34 RRTs, 14 TBR/TBLs, 6 GALS, 34 β-AGP:GALTs, 10 GALT29A/GALT31A, 2 RAY1s, 4 FUTs, and 16 GlcATs. The PCC result showed that there was a high correlation with 1 RG-I:GalAT, 1 RRT, 2 GALSs, 3 AGP:GALTs, and 2 TBRs (Fig. 9D; Supplemental Table S8F). Similar way PCC analysis between 5 RG-I β-1,6-linked galactan epitopes (content goes up during 6-25 DPA) and the same transcripts mentioned above revealed that 1 RG-I:GalAT, 2 RRT, 2 AGP:GALTs, 2 GlcATs and 2 TBL3s were highly correlated (Fig. 9E; Supplemental Table S8G). The same transcripts were PCC analyzed with 7 arabinogalactan (AG/AGP) epitopes and it showed that 1 RG-I:GalAT, 1 RRT, 2 GALSs, 3 AGP:GALTs, and 1 TBR were highly correlated (Fig. 9F; Supplemental Table S8H).

#### iii) Correlation analysis of xylan related epitopes and the corresponding transcripts

The xylan β-1,4-xylosyltransferases (IRXs; irregular xylem), UDP-GlcA: xylan α-glucuronyltransferases (GUXs), glucuronoxylan methyltransferase-like proteins (GXMT), and xylan arabinosyltransferases (XATs) are responsible for the synthesis of xylan molecules (Fig. 2). IRXs are involved in xylan backbone synthesis. GUXs are responsible for adding α-1,2-d-glucuronic acid (GlcA) to the xylan backbone and the GXMTs methylate GlcA molecules. XATs are involved in decorating xylan backbones with arabinose, which results in arabinoxylans (Smith et al. 2017). Several O-acetyltransferases (ESK/RWA/TBL) are required to acetylate the xylan backbone. It was found that cotton fiber has 22 IRXs, 16 GUXs, 30 GXMTs, and 40 O-acetyltransferases (ESK/RWA/TBL) homologous to *Arabidopsis* genes including from both A and D cotton genomes (Supplemental Table S8B).

The profiles of the five of the xylan backbone (Xyl-BB) epitopes from the pectin-enriched fraction were PCC analyzed with the transcript profiles of 22 IRXs and 40 O-acetyltransferases (ESK/RWA/TBL), which are the enzymes involved in backbone synthesis. The results showed that 2 IRXs, 2 ESKs, and 1 TBL30 correlated significantly with these epitopes (Fig. 10A; Supplemental Table S8I). PCC analysis of Xyl-GlcA epitope profile with the transcript profiles of 22 IRXs, 16 GUXs, and 40 ESKs/RWAs/TBLs showed that 2 IRXs, 2 GUXs, 2 ESKs, and 2 TBL30s correlated significantly with this epitope (Fig. 10B; Supplemental Table S8J).

**Figure 10.**
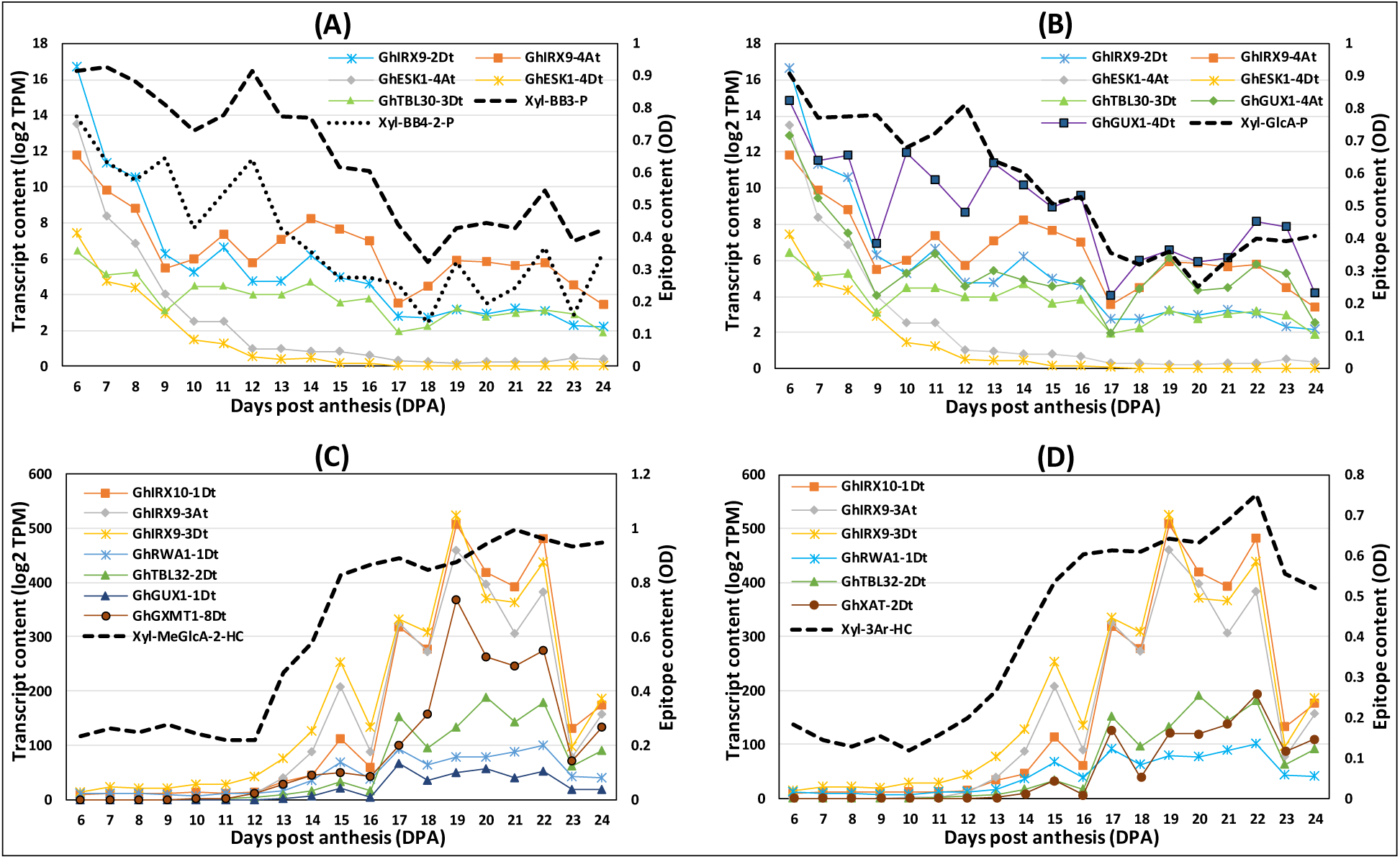
Profiles of xylan epitopes and the correlated profiles of the transcripts of enzymes involved in synthesizing the epitopes. **A)** Profile of xylan backbone epitopes (Xyl-BB3-P and Xyl-BB4-2-P) and correlated representative transcripts. **B)** Profile of glucuronoxylan epitope (Xyl-GlcA-P) and correlated representative transcripts. **C)** Profile of methylated glucuronoxylan epitope (Xyl-MeGlcA-HC) and correlated representative transcripts. **D)** Profile of arabinoxylan epitope (Xyl-3Ar-HC) and correlated representative transcripts. Refer to Supplemental Table S8 (8B, 8I to 8L) for the full list of correlated and non-correlated transcripts. Refer to Fig. 2 for the details of epitope structures and the CW synthesizing enzymes involved.

Interestingly, the PCC analysis of the methylated version of Xyl-GlcA (Xyl-MeGlcA-2-HC) epitope revealed many significantly correlated transcripts, which include 10 IRXs, 5 GUXs, 8 GXMTs, 2 ESKs, 3 RWAs, and 6 TBLs (Fig. 10C; Supplemental Table S8K). PCC analysis was performed between Xyl-3Ar epitope from hemicellulose-enriched fraction and the corresponding transcripts (10 XATs, 22 IRXs, and 40 ESK/RWA/TBL) (Fig. 10D). Obtained results revealed that 12 IRXs, 2 ESKs, 5 RWAs, 6 TBLs, and 1 XAT were highly correlated with the Xyl-3Ar-HC epitope (Fig. 10D; Supplemental Table S8L).

#### iv) Correlation analysis of callose epitope and the corresponding transcripts

Callose is a β-1,3-linked glucan (linear and branched) synthesized by glucan synthase-like enzymes (GSLs). *Arabidopsis* has 8 GSLs (Feng et al. 2021). In cotton fiber, 32 GSLs from A and D genomes were expressed (Supplemental Table S8B). PCC analysis of the callose epitope profile (Cal-P) with the transcript profiles of 32 GSLs showed that 3 GSLs significantly correlated (Fig. 11A; Supplemental Table S8M).

**Figure 11.**
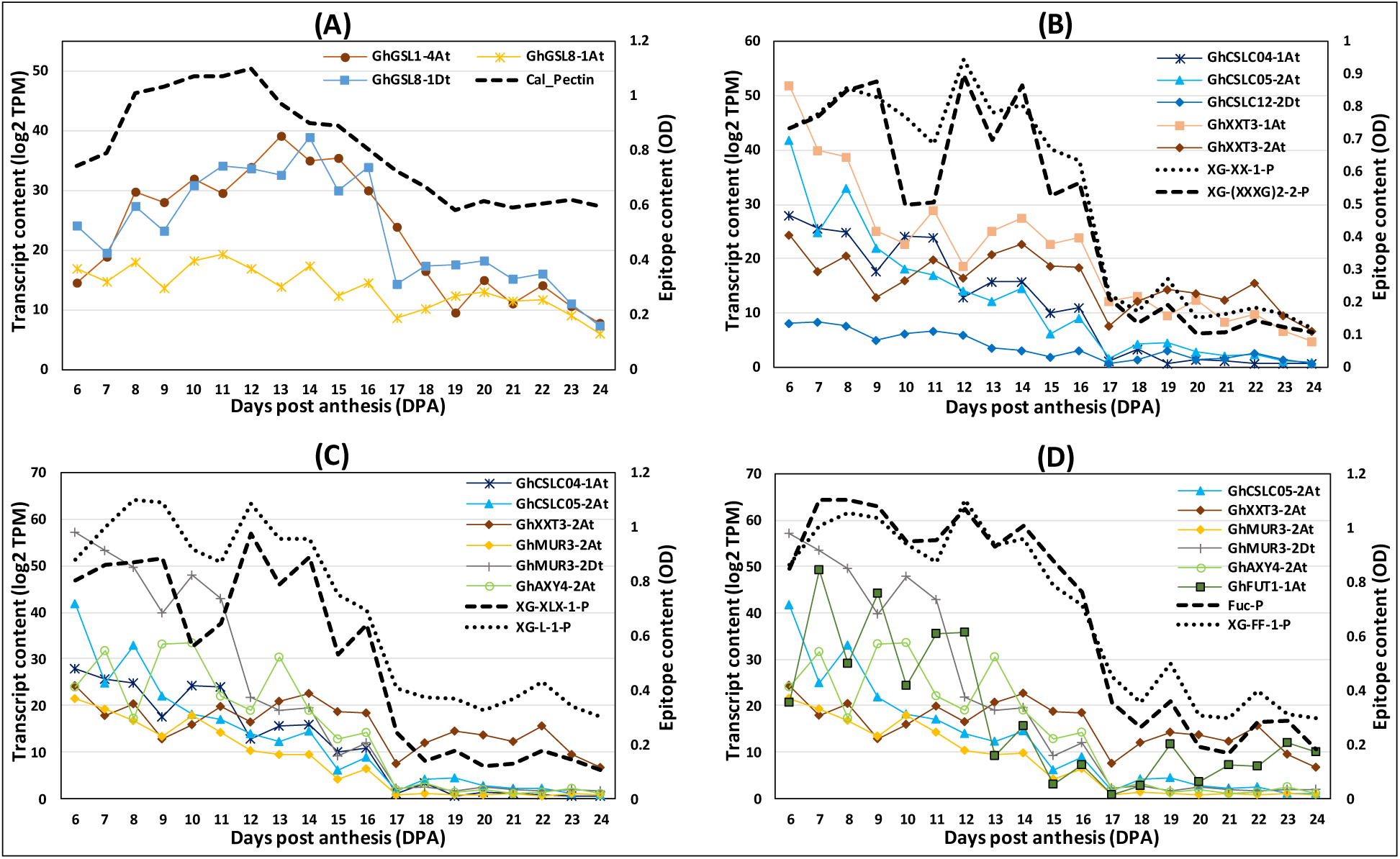
Profiles of callose / xyloglucan epitopes and the correlated profiles of the transcripts of enzymes involved in synthesizing the epitopes. A) Profile of callose epitope (Callose-P) and correlated representative transcripts. B) Profile of xylosylated xyloglucan epitopes (XG-XX-1-P and XG-(XXXG)2-2-P) and correlated representative transcripts. C) Profile of galactosylated xyloglucan epitopes (XG-XLX-1-P and XG-L-1-P) and correlated representative transcripts. D) Profile of fucosylated xyloglucan epitopes (Fuc-P and XG-FF-1-P) and correlated representative transcripts. Refer to Supplemental Table S8 (8B, 8M to 8P) for the full list of correlated and non-correlated transcripts. Refer to Fig. 2 for the details of epitope structures and the CW synthesizing enzymes involved.

#### v) Correlation analysis of xyloglucan related epitope and the corresponding transcripts

In *Arabidopsis*, xyloglucan β-1,4-glucan synthases (cellulose synthase-like-C enzymes; CSLCs; CSLC4/5/6/8/12), UDP-Xyl: xyloglucan α-1,6-xylosyltransferases (XXTs), xyloglucan β-1,2-galactosyltransferases (MUR3 and XLT2) and a GDP-L-Fuc: α-1,2-L-fucosyltransferases (FUT1) are involved in biosynthesis of xyloglucans (XyG) (Fig. 2) (Julian and Zabotina 2022). The CSCLs are involved in the synthesis of the glucan backbone of XyG. Five different XXTs (XXT1 through XXT5) attach xylose residues to the glucoses at different positions in the glucan backbone. Further, MUR3 and XLT2 galactosylate the specific xyloses in XyG. Finally, FUT1 fucosylates the galactose in the second side chain of XyG subunit. In addition, O-acetyltransferases (AXY4/9) are known to acetylate the galactose residues in the second side chain of XyG. The search for genes homologous to *Arabidopsis* genes showed that cotton has 16 CSLCs, 10 XXTs, 8 MUR3s, 4 XLT2s, 4 FUT1s, and 12 O-acetyltransferases (AXY4/9), including from A and D cotton genomes (Supplemental Table S8B).

PCC analysis of 4 xylosylated glucan backbone epitopes of XyG (XG-XXs and XG-XXXGs) from the pectin-enriched fraction 16 CLSCs and 10 XXTs, showed that 7 CLSCs and 5 XXTs were highly correlated (Fig. 11B; Supplemental Table S8N). In another set of PCC analyses, the profiles of 5 galactosylated XyG (XG-XLXs and XG-Ls) epitopes significantly correlated with the transcripts of 6 CLSCs, 5 XXTs, 5 galactosyltransferases (4 MUR3 and 1 XLT2), and 1 AXY4 (Fig. 11C; Supplemental Table S8O). PCC analysis of 3 fucosylated XyG (Fuc and XG-FF) epitopes and the transcripts of enzymes putatively involved in synthesizing these epitopes showed a high correlation for 11 CLSCs, 5 XXTs, 5 galactosyltransferases (4 MUR3 and 1 XLT2), 1 FUT1, and 1 AXY4 (Fig. 11D; Supplemental Table S8P).

## Discussion

Cotton fiber CW polysaccharide composition and complex remodeling of polysaccharide epitopes during development ultimately determines the outcome of fiber quality. Significant efforts have been made to study the cotton fiber CW polysaccharide compositional changes during development (Meinert and Delmer 1977; Singh et al. 2009; Avci et al. 2012; Hernandez-Gomez et al. 2017; Guo et al. 2019; Pettolino et al. 2022). However, the previous studies were carried out on a limited scale, included only a few DPAs, and, as a result, did not reveal finer nuances and dynamics of the spectrum of CW polysaccharide epitopes during development. The uniqueness of the present study is that the glycome, transcriptome, and fiber-phenotypes profiling was carried out using cotton bolls collected daily for 20 consecutive days (6 to 25 DPA). The 20 days include key phases of fiber development, where extensive multiple phases of CW remodeling occur that determine the fiber quality. This large-scale approach allowed us to discover dynamic changes the occur in fiber CW during development with better resolution.

### Predicted polysaccharide epitopes and CW synthesizing enzymes potentially contributing to fiber elongation

The glycome profile data revealed that most of the highly branched RG-I pectin epitopes, like β-1,4-linked galactans, β-1,6-linked galactans, and arabinogalactans were present at high amounts at earlier stages of fiber development (Figs. 4 and 6; Tables 1 and 3) and significantly decreased around 16-18 DPA, coinciding with the CFML degradation (Guo et al. 2019) and cessation of fiber elongation (Howell et al. 2024 unpublished). A similar profile was observed for buffer-soluble callose, xyloglucans, and xylans in our and earlier studies (Singh et al. 2009; Avci et al. 2012; Guo et al. 2019). The polysaccharide content estimation also clearly showed the reduction of pectins and hemicelluloses percentages in CW during the transition stage, which is also corroborated by the decrease in monosaccharides like galacturonic acid, xylose, galactose, and fucose (Fig. 1; Supplemental Tables S1 and S2). Only glucose content was found to increase in both the buffer and alkali-soluble extracts, which might be due to the soluble amorphous cellulose leached out from the CW during extraction.

It is known that the large side chains in highly branched pectins prevent extensive interactions among adjacent polysaccharides to maintain plasticity to support cell elongation, whereas less branched polysaccharides promote bonding and rigidification of CW, thus preventing cell elongation (Atmodjo et al. 2013). For example, highly branched RG-I (with arabinan and galactan side chains) has reduced interaction with cellulose or xyloglucans as a result of steric impediments, which resulted in increased CW plasticity (Zykwinska et al. 2005; Hofte and Voxeur 2017). Also, the highly branched RG-I was reported to prevent the formation of cross-linking of HG by Ca2+ bridges (Jones et al. 2003). Increased methyl-deesterification of HG was reported to increase the cross-linking of HG domains by Ca2+ bridges and contributing to CW stiffening (Kaczmarska et al. 2022). Our and also earlier study (Pettolino et al. 2022) demonstrated that the content of alkali-soluble methyl-esterified-HG in fiber decreased after 13 DPA, whereas the deesterified-HG remains high (Tables 1 and 4). These results provide further proof that decrease in branched RG-I, combined with an increase in deesterified HG and cellulose content at the beginning of the transition phase, might be contributing to rigidification of the CW and hindering further cell elongation.

Using correlation data, we are able to predict a subset of potential glycosyltransferases that are likely to synthesize HG. The PCC analysis demonstrated that the expression of 3 GAUT1, 1 GAUT3, 2 GAUT4, 2 GAUT6, 1 GAUT9, 1 GAUT10, 1 GALT3, and 2 GALT9 significantly correlated with the HG epitope profiles (Fig. 9A; Supplemental Table S8C). The significantly correlated cotton homologous genes identified in our study appear to differ from the key GAUTs found in *Arabidopsis*. In *Arabidopsis*, GAUT1 is the only essential catalytically active enzyme that forms obligatory complexes with GAUT7 (Atmodjo et al. 2013; Engle et al. 2022). Since we did not observe a similar correlation for cotton homolog to GAUT7, it will be interesting to investigate in the future whether the cotton GAUT1 protein also gets processed and requires any other GAUT protein to be localized in cotton Golgi.

Similar to methyl esterified-HG, the content of buffer-extractable callose decreased after 13 DPA (Fig. 11A). In an earlier study, the presence of two types of callose in cotton fiber was reported, the buffer soluble highly branched callose and less soluble unbranched callose that can be extracted only in strongly basic conditions (Pettolino et al. 2022). It is possible that loosely bound buffer-soluble branched callose present in PCW also contributes to the plasticity and elongation of the fiber.

A companion study reported that the growth rate of fiber decreases during the entire developmental time course, and the turgor pressure of the fiber peaks at around 16 DPA and decreases rapidly to reach a low level at 22 DPA, coinciding with the complete cessation of fiber elongation (Howell et al. 2024 unpublished). As reported previously, the turgor pressure plays a role in the elongation of fiber and reaches its peak coinciding with the attainment of maximum fiber length (Howell et al. 2024 unpublished). Profiles of many of the highly branched RG-I epitopes significantly correlated to the fiber growth rate and the estimated turgor pressure profile (Figs. 4 and 8; Supplemental Table S7). The buffer-soluble xyloglucans and xylans similar to pectins began to decrease as early as at 12 DPA, and the profile of these epitopes also strongly correlated with the fiber growth rate curve (Figs. 4 and 8; Supplemental Table S7). Most likely, presence of these epitopes in the CW supports CW plasticity and building turgor pressure to promote fiber elongation. On the other hand, degradation of these highly branched epitopes combined with an increase in SCW cellulose accumulation (Figs. 1A and 4) might facilitate the CW rigidification, reduction in turgor pressure, and cessation of fiber elongation.

RG-I primary backbone is decorated with the secondary β-1,4-galactan backbone, arabinans, β-1,6-galactans, and arabinogalactans. A sharp decline of β-1,4-galactans immediately after 16 DPA, shown in our study (Fig. 4B), has also been reported earlier, coinciding with the cessation of fiber elongation (Singh et al. 2009; Guo et al. 2019). These authors proposed that β-1,4-galactan molecules may be components of CFML and correlated with fiber elongation. An *Arabidopsis* GALS protein (GALS1), which participates in β-1,4-galactan secondary backbone synthesis, was functionally characterized by biochemical assays (Liwanag et al. 2012). We found two of the cotton GALS1, homologous to *Arabidopsis* genes, significantly correlated with the β-1,4-galactan epitope profiles (Fig. 9C; Supplemental Table S8E). An earlier transcriptomic and glycomic study in three cotton species (*G. hirsutum*, *G. barbadense* and *G. arboretum*) also predicted the same GALS1 homolog gene to be involved in β-1,4-galactan synthesis (Guo et al. 2019).

The buffer extractable xyloglucan epitopes present at high content at earlier DPAs significantly decreased during the transition stage (Fig. 4B). In *Arabidopsis*, the completely branched structure of xyloglucan (XLFG type) is synthesized by CLSCs (CSLC4/5/6/8/12), XXTs (XXT1/2/3/4/5), galactosyltransferases (MUR3/XLT2), and a fucosyltransferase (FUT1) (Pauly and Keegstra 2016; Julian and Zabotina 2022; Zhang et al. 2023). From our transcriptome analysis, we did not find cotton homologous genes for CSLC6, CSLC8, XXT1, XXT4, and XXT5 (Supplemental Table S8B). The PCC analysis for the CSLC transcripts revealed that transcripts of two genes, each from CSLC4, CSLC5, and CSLC12 highly correlated with all the XyG epitopes (Fig. 11; Supplemental Tables S8, N to P). Earlier studies in *Arabidopsis* reported all five CSLCs are involved in the XyG synthesis (Kim et al. 2020), however, to different extent in different tissues. The authors demonstrated that CSLC4 and CSLC5 had the highest levels of expression in vegetative tissues, whereas expression of CSLC6 and CSLC12 was specific to flowers and seeds.

Among cotton XXTs, one homologous to *Arabidopsis* XXT2 and four to XXT3 were found to significantly correlate with the XyG epitopes (Fig. 11; Supplemental Table S8, N to P). In *Arabidopsis*, XXT1, XXT2, and XXT5 have the highest levels of expression in most tissues (Chou et al. 2012; Zabotina, 2012) and were demonstrated to be the major players in XyG synthesis (Zabotina et al. 2008; Zabotina et al. 2012). In *Arabidopsis,* XXT5 plays a dominant role, and XXT3 and XXT4 play a minor complementary role in adding xylosyl residue specifically to the third glucose in the glucan backbone to synthesize XXXG-type XyGs (Culbertson et al. 2018; Zhang et al. 2023). On the contrary, our correlation data suggest that in cotton fiber, XXT2 and XXT3 are the key enzymes involved in XyG synthesis. It is plausible that in cotton fiber, four of the XXT3 proteins might be substituting the function carried by XXT5 in *Arabidopsis*. However, in the vegetative tissues, the expression of the XXT genes might vary, and it will require separate investigation.

Correlation analysis revealed that the transcripts of four MUR3, one XLT2, one fucosyltransferase, and an O-acetyltransferase significantly correlated with the XyG epitopes (Fig. 11, C and D; Supplemental Table S8, O and P). A previous study using small-scale transcriptomic and glycomic analyses in cotton fiber of two species reported the possible involvement of CSLC4, XXT2, MUR3, and FUT1 in cotton fiber xyloglucan synthesis, which corroborates our findings (Guo et al. 2019). In our large-scale study, we found several additional correlating genes. For example, transcripts of four homologous MUR3 genes were found to highly correlate with XyG glycome profiles (Fig. 11; Supplemental Table S8, O and P). However, the real contribution of all of these enzymes in XyG biosynthesis during fiber development will need further experimental validation.

In previous studies of cotton fiber, a lot of attention was paid to CFML, an adhesive layer of fiber CW that supports fiber elongation and its degradation leads to cessation of fiber elongation (Singh et al. 2009; Hernandez-Gomez et al. 2017). Pectins are known to support cell-cell adhesion (Kaczmarska et al. 2022). Previous immunolabeling-TEM studies reported the presence of HGs, and xyloglucans, and possibly RG-I-galactan epitopes in CFML and proposed that these particular polysaccharides contribute to fiber elongation (Singh et al. 2009; Avci et al. 2012; Guo et al. 2019). Our study clearly showed that HG, xyloglucans, and also many specific branched RGI epitopes (galactans, arabinogalactans) significantly reduced during the transition stage coinciding with CFML degradation (Tables 1 and 3). Taken together, our results and earlier studies indicate that the most likely combination of the HGs, xyloglucans and branched RG-Is present in the CFML and other polysaccharides and CW loosening proteins, such as expansins present at PCW might collectively control the fiber plasticity and elongation.

### Predicted polysaccharide epitopes and CW synthesizing enzymes potentially contributing to fiber strength

Events such as CFML degradation, accumulation of callose, and SCW cellulose leads to fiber CW rigidification and increased strength (Maltby et al. 1979; Jaquet et al. 1982; Rowland et al. 1984; Martel and Giovannoni 2007; Vicente et al. 2007; Pettolino et al. 2022; Fang et al. 2023). Interestingly, few specific polysaccharide epitopes (Xyl-MeGlcA-1-HC, Xyl-MeGlcA-2-HC, Xyl-3Ar-HC, RG-I-Gal1/2/3-β-1,6-galactans, AG-3-2-HC, AG-4-HC, and AG-4/Gal3-BB-HC) were absent or low at earlier DPAs but increased significantly between 12-14 DPAs, which coincides with the beginning of transition stage and SCW cellulose synthesis (Fig. 5; Table 2). Our correlation analysis showed a large number of xylan β-1,4-xylosyltransferases (IRX9, 10, 14), O-acetyltransferases (EKS1/RWAs/TBLs), xylan α-glucuronyltransferases (GUXs), and glucuronoxylan methyltransferases (GXMTs) highly correlated with the Xyl-MeGlcA and Xyl-3Ar epitopes profile (Fig. 10, C and D). These results indicate the prime importance of the potential involvement of these xylan epitopes and these large sets of enzymes in SCW assembly and fiber-strengthening process.

An earlier study using four domesticated species of cotton also reported a heteroxylan epitope and few correlated transcripts (Hernandez-Gomez et al. 2015) increased during transition stages, which matches to the findings of our study. The authors proposed that the heteroxylan epitope might be involved in the regulation of cellulose deposition in SCW. Interestingly, it was shown earlier that glucuronoxylans (Xyl-GlcA) are the predominant type of xylan found within the SCW of hardwoods and herbaceous dicot species like poplar and *Arabidopsis* (Jacobs et al. 2001; Zhong et al. 2005; Mazumder et al. 2012; Smith et al. 2017).

Previous experiments showed that unbranched xylan backbone is structurally similar to cellulose and can spontaneously bind to cellulose microfibrils affecting their orientation and aggregation during CW synthesis (Vian et al. 1986; Atalla et al. 1993; Smith et al. 2017). Studies in woody tissues showed that glucuronoxylans deposited in the transition zones between the “S” layers participate in organizing the cellulose microfibrils and change their orientation (Vian et al. 1986; Atalla et al. 1993). It is worth noting that in our study, the profile of the three xylan epitopes and the correlated transcripts highly matches the profile of cellulose content, the expression of SCW synthesizing CESAs, and the microfibril orientation (Figs. 1A, 8C, 8D, 10C and 10D; Supplemental Fig. S3) and the CW thickness phenotype profile reported earlier in a collaborative study (Howell et al. 2024 unpublished). The cellulose content and highly correlated SCW-CESAs (CESAs 4/7/8) were found to increase dramatically around 16 DPA, which coincides with the transition stage and the beginning of SCW accumulation (Supplemental Fig. S3; Supplemental Table S9). The highly correlated SCW-CESAs identified in cotton matches to the known homologous SCW-CESAs present in *Arabidopsis* (CESAs 4/7/8) (Kim et al. 2019).

Interestingly, few of the RG-I-β-1,6-galactan epitopes and significantly correlated transcripts that goes up during SCW accumulation (Table 2; Fig. 5) highly matches the three xylan epitopes and microfibril orientation profiles (Fig. 8C; Supplemental Table S7). In the future, it might be worth investigating whether any of the three xylan epitopes (Xyl-MeGlcA-1-HC, Xyl-MeGlcA-2-HC, Xyl-3Ar-HC) and RG-I-β-1,6-galactan epitopes identified here participate in cellulose microfibril re-arrangement, thus promoting SCW synthesis and increased strength of the mature fiber. The involvement of callose and cellulose in “winding CW layer” formation and its contribution to the strengthening of the fiber has been proposed as well (Kerr, 1946, Hsieh et al. 1995; Hinchliffe et al. 2011).

In conclusion, this paper integrated glycome profiling, transcriptomics, and multiple phenotypes with an unprecedented temporal resolution. The fiber growth process is not simple, and there is a complicated temporal sequence of CW polysaccharide remodeling that defines morphological and material property traits that underlie fiber quality. Our study uncovered some of the highly branched RG-I pectin epitopes, in addition to buffer-soluble HGs and xyloglucans, could play an important role in the formation of the CFML and PCW and contribute to fiber plasticity and elongation. On the other hand, the identified heteroxylans (Xyl-MeGlcA-1-HC, Xyl-MeGlcA-2-HC, Xyl-3Ar-HC) might function as a regulating factor for SCW cellulose microfibrils arrangement and further strengthening of the fibers. Our study also identified several highly promising correlated cotton polysaccharide-synthesizing enzymes. In the future, additional molecular and biochemical experiments will be required to validate the biochemical role of the identified polysaccharide epitopes and putative enzymes in cotton fiber elongation and strength.

## Materials and Methods

### Cotton plant growing conditions and boll collection

*Gossypium hirsutum* accession TM1 plants were grown in a growth chamber (Conviron E-15, Controlled Environments Inc. ND, USA) under controlled conditions. Plants were grown individually in a two-gallon pot containing a potting mix of 4:2:2:1 ratio of soil:perlite:bark:chicken grit. Growth chambers were programmed for 16-hour days with 500 µmol of light and a temperature of 28° C. At anthesis, flowers were self-pollinated and tagged. At each daily developmental timepoint from 6 to 25 DPA, three replicate bolls were collected at mid-day, labelled, and stored at -80° C until used for CW or RNA extraction.

### Cell wall and polysaccharide extraction

Alcohol-insoluble CW material and, further, the pectin- and hemicellulose-enriched polysaccharide fractions from the cotton fiber were extracted as per an established protocol (Avci et al. 2012). In brief, using a sharp razor blade, fibers were removed from the seeds. The harvested cotton fibers were ground in liquid nitrogen, and from the powdered fiber the CW was extracted by using different solvents. From the CW material, polysaccharides were fractionated by sequential extraction using two different solutions, first using a 50mM CDTA: 50mM ammonium oxalate (1:1) buffer, followed by 4M KOH to extract pectin- and hemicellulose-enriched polysaccharides, respectively (Supplemental Fig. S1) (Zabotina et al. 2012). The final pellet (containing a mixture of amorphous and crystalline celluloses) that remained after the 4M KOH extraction was also weighed. The crystalline cellulose content present in this final pellet was measured by the Updegraff reagent method (Updegraff, 1969).

### Glycome profiling of epitopes of pectin-enriched and hemicellulose-enriched polysaccharide fractions

The amount of sugar in the pectin- and hemicellulose-enriched fractions from each sample was determined by the phenol-sulfuric acid method (Pattathil et al. 2012), and each well of ELISA plate (Costar 3595) was loaded with an equal amount of polysaccharide sample dissolved in water (50 µl/well from a 60 µg/µl solution). Glycome profiling is an ELISA-based method and it was carried out as described previously (Pattathil et al. 2010, 2012; Zabotina et al. 2012) with 71 different polysaccharide epitope-specific antibodies (Supplemental Table S3) that were selected based on the literature (Avci et al. 2012; Thorne et al. 2023).

Monosaccharide composition was determined according to previously described methods (Brenner et al. 2012). The CW fractions (1 mg) were hydrolyzed with 2M trifluoroacetic acid and analyzed by high-performance liquid chromatography using an anion-exchange chromatography column (Dionex, Sunnyvale, CA). The column was calibrated using nine monosaccharide standards (Brenner et al. 2012).

### Self-organizing maps (SOM) clustering

SOM is a machine learning method for clustering analysis (Kohonen 1990; Wehrens and Kruisselbrink 2018). We used the SOM algorithm to cluster the glycome epitopes from two separate interpolated datasets: pectin-enriched and hemicellulose-enriched polysaccharide fractions. The SOM grid structures were adopted as a grid structure with 5 rows and 4 columns for each data set.

### RNA extraction and RNA sequencing

In parallel to glycome profiling, RNAseq and transcriptomic profiling were carried out for the cotton fibers collected simultaneously for the same number of days (6 to 25 DPA). Briefly, a modified version of the Sigma Plant Spectrum Total RNA kit (Sigma-Aldrich) was used to extract RNA from the fibers. Library construction (NEBNext Ultra II RNA Library Prep Kit) and sequencing (as PE150; Illumina NovaSeq 6000) were conducted by the Iowa State University DNA facility. Data were processed using Trimmomatic version 0.39 (trimmomatic/0.39-da5npsr) (Bolger et al. 2014) from Spack (Gamblin et al. 2015) for read and quality trimming. The reference transcriptome was a species-specific, homoeolog-diagnostic reference transcriptome based on the *G. raimondii* (Paterson et al. 2012) genome annotation. The transcripts per million (TPM) output by Kallisto for each sample were imported into R/4.2.2 (R Core Team 2022) and RNA-seq quality was assessed by consistency of number of genes expressed over time and among replicates. The number of expressed genes per sample (TPM > 0) was plotted across developmental time using ggplot2, and visual outliers were discarded. RNAseq reads were deposited into the NCBI-SRA under PRJNA1099209.

### Statistical analysis

All experimental mean data, including Pearson correlation coefficient (PCC) analysis, were subjected to statistical analysis using R software. The data of each experiment were initially tested for significance by analysis of variance (ANOVA), and subsequently, Fisher’s protected least significant difference (LSD) test value was used to compare treatment means at *p* < 0.05.

### Accession numbers

RNAseq reads of cotton fibers collected from 6 to 25 DPA were deposited into the NCBI-SRA under PRJNA1099209.

## Acknowledgements

This study was supported by the National Science Foundation (NSF) - Plant Genome Research Project (Grant No. 1951819).

## Competing interests

None declared.

## Author contributions

DBZ, JFW, OAZ, and JX conceived the research. JJJ, and AGL grew the cotton plants, pollinated, collected bolls, and extracted RNA from fibers. CEG and JFW processed and analyzed the RNAseq transcriptome data. SS, ASM, and LES extracted polysaccharides from cotton fibers. SS and OAZ conducted glycome profiling and monosaccharide analysis of polysaccharides and analyzed the data. PY and JX generated SOM and conducted PCC analysis. YL generated heat map. YL, ELM, and DBS conducted fiber phenotype measurements and analyzed the data. SS wrote the article, with significant input from OAZ, JFW, DBS, YL and CEG. All authors contributed with comments and suggestions to finalize the manuscript.

## Supplemental data

**Supplemental Figure S1.** Schematic representation of cotton cell wall polysaccharide extraction and analysis.

**Supplemental Figure S2.** Pearson correlation coefficient (PCC) distribution of the correlated polysaccharide epitopes and transcripts.

**Supplemental Figure S3.** Correlation analysis of profiles of cotton fiber cellulose content and the transcripts of cellulose synthase (CESA) enzymes involved in primary and secondary cell wall synthesis.

**Supplemental Table S1.** Weight and percentage content of cotton fiber cell wall polysaccharides (6 - 25 DPA).

**Supplemental Table S2.** Monosaccharide content of polysaccharide fractions of cotton fiber (6 - 25 DPA).

**Supplemental Table S3.** List of polysaccharide specific antibodies used in this study and their binding epitopes.

**Supplemental Table S4.** Glycome profiling data of pectin-enriched fractions of cotton fiber (6 - 25 DPA).

**Supplemental Table S5.** Glycome profiling data of hemicellulose-enriched fractions of cotton fiber (6 - 25 DPA).

**Supplemental Table S6.** Self-organizing map (SOM) groups of glycome profiled polysaccharide fractions and the epitopes present in each group.

**Supplemental Table S7.** Correlation analysis of cotton fiber phenotypes and the polysaccharide epitopes.

**Supplemental Table S8.** List of cell wall polysaccharide synthesizing glycosyl transferases and the correlation analysis with their corresponding polysaccharide epitopes.

**Supplemental Table S9.** Correlation analysis of cellulose and cellulose synthases (CESAs) of *Gossypium hirsutum*.

## Notes

### Competing Interest Statement

The authors have declared no competing interest.

