## Supplemental Figure S1 for "Daily glycome and transcriptome profiling reveals polysaccharide structures and glycosyltransferases critical for cotton fiber growth"

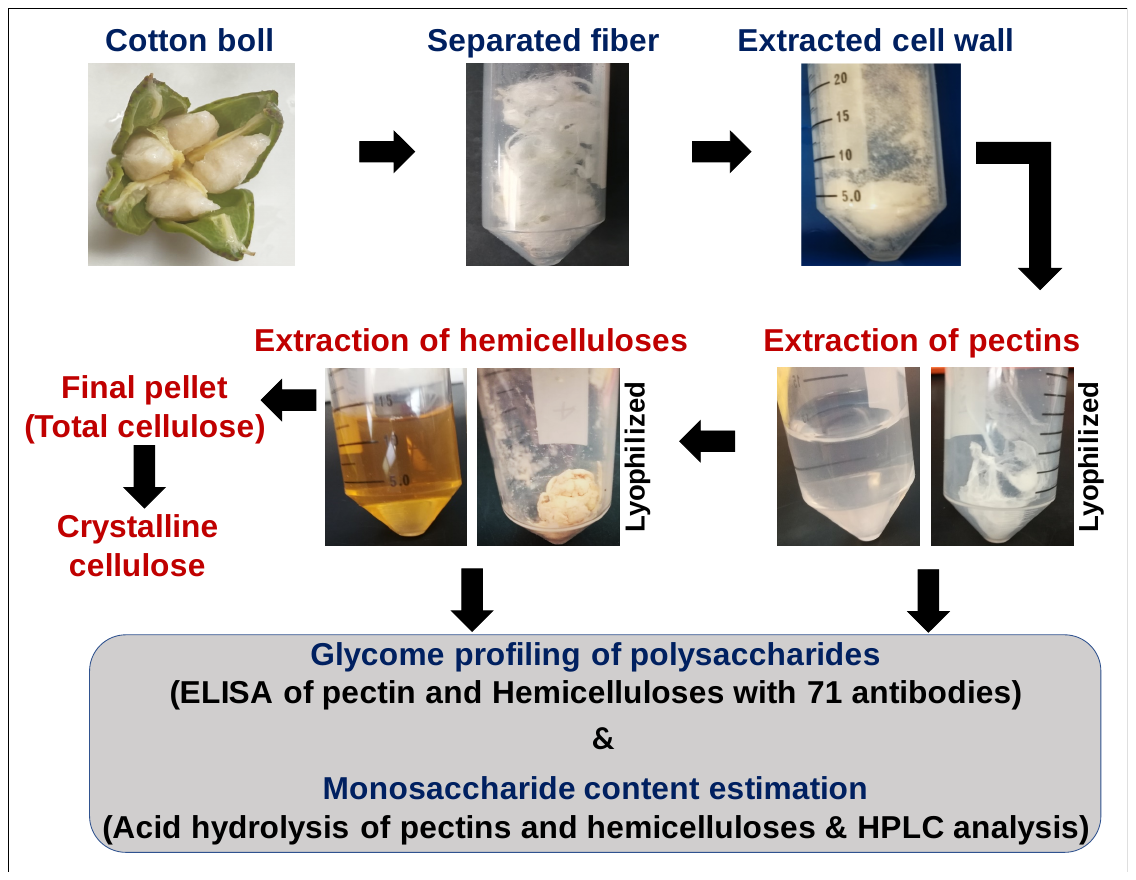

**Supplemental Figure S1.** Schematic representation of cotton cell wall polysaccharide extraction and analysis. The cell wall (CW), pectin, and hemicellulose fractions from three biological replications of 6 to 25 DPA cotton fibers were extracted as described previously (Zabotina et al. 2012). Briefly, the fibers were separated from the seeds, ground to a fine powder in liquid nitrogen, and the CW was extracted by using successive organic solvents. From the CW, pectin and hemicellulose polysaccharides were extracted successively by using 50 mM CDTA:50 mM ammonium oxalate (1:1) buffer (*v*:*v*) and 4 M KOH, respectively. The extracts were dialyzed, and dried by lyophilization. The final pellet left out after pectin and hemicellulose extraction has a mixture of both amorphous and crystalline celluloses. From the final pellet, the crystalline cellulose content was estimated by Updegraff method (Updegraff 1969). Glycome profiling of the pectin-enriched and hemicellulose-enriched extracts was performed as previously described (Pattathil et al. 2012) using 71 different antibodies. Monosaccharide composition analysis was determined according to previously described methods ([Brenner et al. 2012](javascript:;)) using high-performance anion-exchange chromatography.
