## Supplemental Figure S2 for "Daily glycome and transcriptome profiling reveals polysaccharide structures and glycosyltransferases critical for cotton fiber growth"

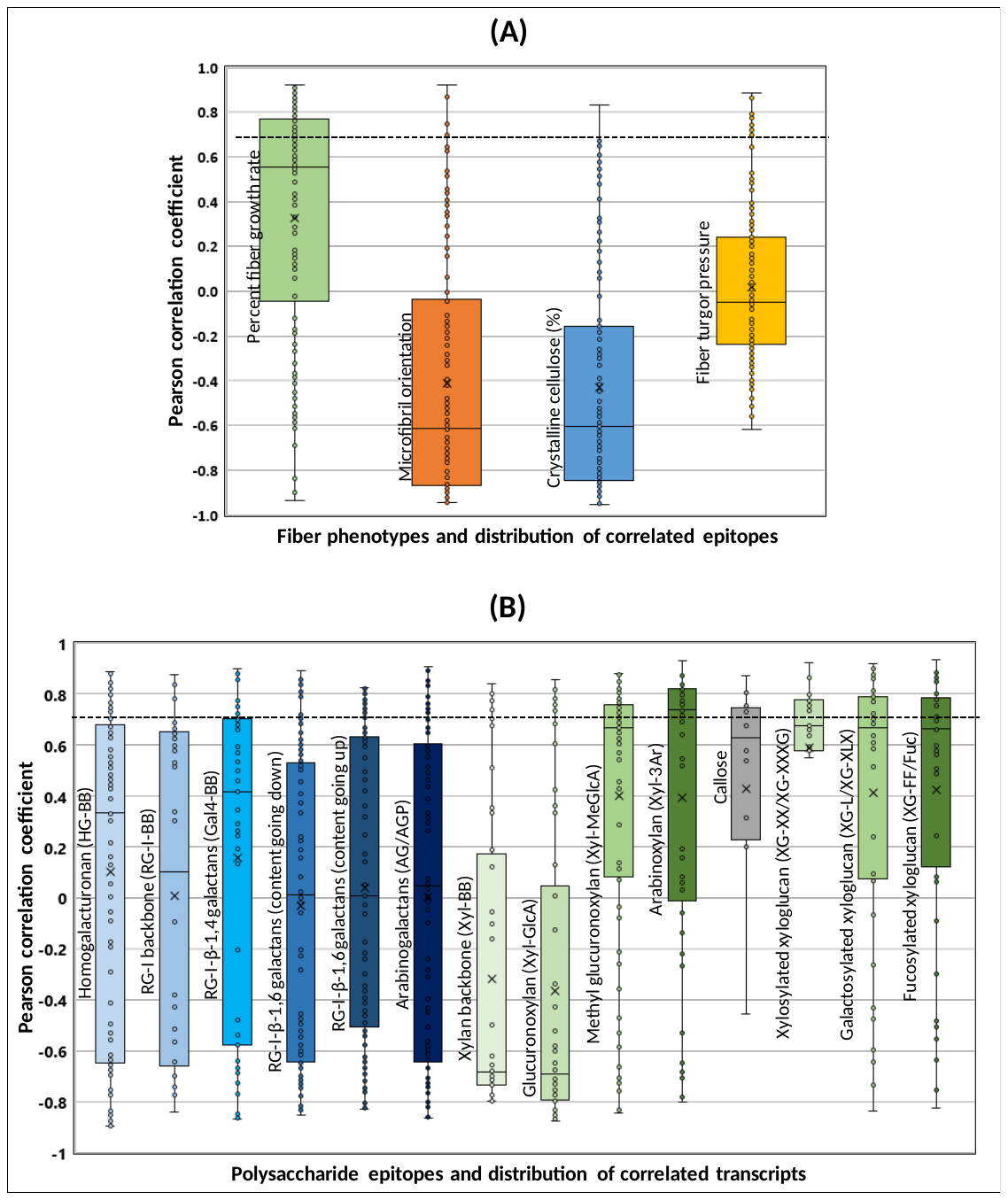

**Supplemental Figure S2.** Pearson correlation coefficient (PCC) distribution of the correlated polysaccharide epitopes and transcripts. **A)** PCC distribution of the polysaccharide epitopes correlated against fiber phenotypes profiles. Epitopes with a PCC value threshold of ≥ 0.7 were considered to be strong and positively correlated with the fiber phenotype profiles. Refer to Supplemental Table 10 for the full list of correlated and non-correlated epitopes and the corresponding fiber phenotypes. **B)** PCC distribution of the transcripts of glycosyl transferase enzymes correlated against the corresponding polysaccharide epitopes. Transcripts with a PCC value threshold of ≥ 0.7 were considered to be strong and positively correlated with the epitope profiles. Refer to Supplemental Table S8 for the full list of correlated and non-correlated transcripts and the corresponding epitopes. The pectin and hemicellulose epitopes profile are color coded by blue and green, respectively.
