## Supplemental Figure S3 for "Daily glycome and transcriptome profiling reveals polysaccharide structures and glycosyltransferases critical for cotton fiber growth"

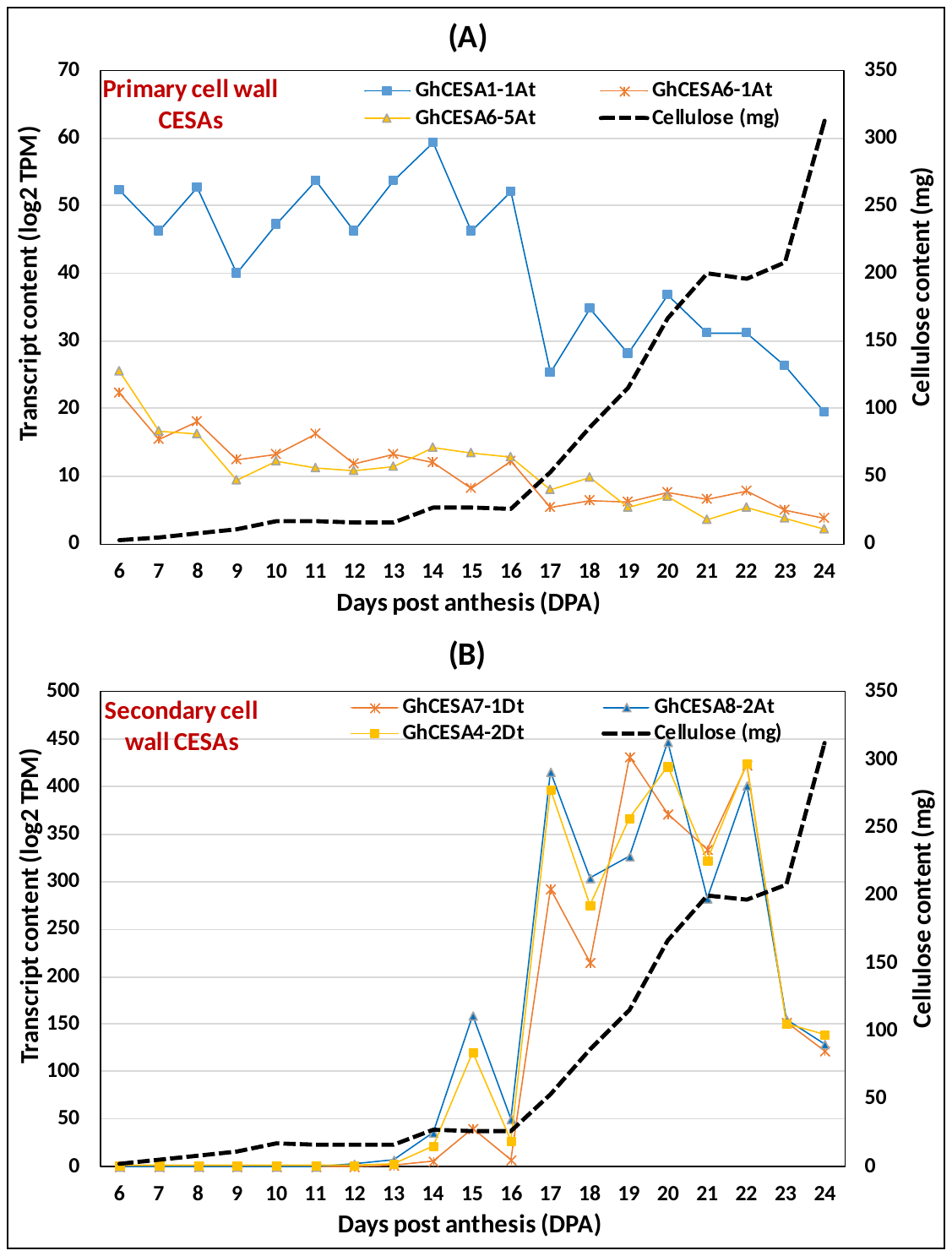

**Supplemental Figure S3.** Correlation analysis of profiles of cotton fiber cellulose content and the transcripts of cellulose synthase (CESA) enzymes involved in primary and secondary cell wall synthesis. **A)** Graph shows bad correlation between the cellulose content and the CESAs involved in primary cell wall (PCW) synthesis. **B)** Graph shows high positive correlation between the cellulose content and the CESAs involved in secondary cell wall (SCW) synthesis. Only few of the representative transcripts are shown here. Refer to Supplemental Table S9 for the full list of correlated and non-correlated transcripts of all the CESAs.
